# Esc1-mediated anchoring regulates telomere clustering in response to metabolic changes

**DOI:** 10.64898/2025.12.09.692953

**Authors:** Myriam Ruault, Isabelle Loïodice, Bradley D. Keister, Antoine Even, Mickaël Garnier, Manuela Baquero-Pérez, David Waterman, James E. Haber, Krastan B. Blagoev, Vittore F. Scolari, Angela Taddei

## Abstract

Spatial organization of yeast telomeres is highly dynamic and regulated by growth conditions. In rich medium, the 32 telomeres group in 3 to 5 foci at the nuclear periphery. This organization is drastically rearranged in long-lived quiescent (Q) cells forming upon carbon source exhaustion: telomere foci assemble into a hypercluster containing most telomeres and located in the center of the nucleus contributing to their long-term viability. Here we explore the mechanisms regulating telomere distribution during this transition. We rule out a modification of telomere-telomere interactions via the telomeric protein Sir3 as the main factor regulating hypercluster formation. However, our physical modeling predicts that telomere anchoring antagonizes telomere clustering. Systematic deletion of all known telomere anchors identifies the inner nuclear membrane-associated protein Esc1, as a key telomeric anchor after the diauxic shift.

Our data support a model where Esc1 mediated anchoring is progressively lost upon entry into quiescence through dephosphorylation of a single residue of Esc1, resulting in the formation of telomere hyperclusters in the center of the nucleus in Q cells.

## Introduction

The three-dimensional organization of the genome is a major regulator of genome functions including DNA repair, transcription, and replication (Bonev & Cavalli, 2016; Li *et al*, 2024; Chiolo *et al*, 2025). In mammals, this organization is highly dynamic, regulated both spatially and temporally throughout the cell cycle and cell differentiation, and influenced by external environmental cues (Dekker *et al*, 2023; Pudelko & Cabianca, 2024). In interphase cells, euchromatin (active chromatin) and heterochromatin (repressive chromatin) are differentially localized within the nucleus, heterochromatin being preferentially associated to the nucleoli and the nuclear periphery, respectively forming NADs (Nucleolus Associated Domains) and LADs (Lamina Associated Domains) (Van Steensel & Belmont, 2017; Allshire & Madhani, 2018; Bizhanova & Kaufman, 2021). When cells leave the cell cycle to enter quiescence or senescence, the 3D architecture of the genome is restructured (Bridger *et al*, 2000; Shaban & Gasser, 2025). Perturbation of chromatin organization in the 3D nuclear space is responsible for multiple diseases such as laminopathies, neurodegenerative diseases and cancer (Manda *et al*, 2023). Numerous molecular mechanisms that affect the spatial organization of chromatin have been described, encompassing chromatin’s physical properties, its compartmentalization, and its interaction with nuclear landmarks such as nuclear envelope proteins, lamina, nuclear pores or the nucleolus (Falk *et al*, 2019; Mirny & Dekker, 2022).

In *S. cerevisiae,* the genome adopts a Rabl-like configuration in which chromosome arms stretch from centromeres, anchored at the spindle pole body to telomeres positioned at the nuclear periphery (Duan *et al*, 2010; Taddei & Gasser, 2012). Although classical markers of heterochromatin typically found in other eukaryotes are missing in yeast, heterochromatin is nonetheless established and maintained in a well-orchestrated manner by the Silent Information Regulator complex (Sir2/Sir3/Sir4) at the 32 telomeres and the cryptic mating-type loci (*HML* and *HMR*) (Gartenberg & Smith, 2016). Although yeast silent chromatin differs from the one of mammalian cells at the molecular level, it is also found enriched at the nuclear periphery. Indeed, the 32 telomeres of a haploid cell are found predominantly at the nuclear periphery where they form 3 to 5 foci (Gotta *et al*, 1996). The local concentration of the SIR protein within these subnuclear compartments favors gene silencing at the nuclear periphery while preventing promiscuous gene silencing elsewhere in the nucleus (Andrulis *et al*, 1998; Taddei *et al*, 2009). This functional organization relies on two conserved features of heterochromatin: homotypic interactions between heterochromatin regions (clustering) and their interaction with nuclear envelope associated proteins.

In yeast, we previously showed that Sir3 is the molecular glue that stick together silent chromatin loci together (Ruault *et al*, 2011, 2021), while telomeres are anchored at the nuclear periphery by multiple redundant pathways involving different mechanisms. One mechanism is mediated through the interaction between telomere-bound proteins such as Sir4, the Yku complex (Hediger *et al*, 2002; Taddei *et al*, 2004), Est1 (Schober *et al*, 2009) and inner-membrane associated proteins such as Esc1 (Andrulis *et al*, 1998, 2002; Taddei *et al*, 2004), Mps3 (Bupp *et al*, 2007; Schober *et al*, 2009), Heh1/Heh2 (Chan *et al*, 2011), Mlp1/Mlp2 (Niepel *et al*, 2013) and Nup170 (Van de Vosse *et al*, 2013; Lapetina *et al*, 2017). Although lamins are not present in yeast, Esc1 may fulfill an analogous structural role (Andrulis *et al*, 2002). Another telomere anchoring mechanism involves the acylation of the telomeric protein Rif1 by Pfa4 resulting in the embedding of the palmitoylated Rif1 in the INM (Park *et al*, 2011).

Nuclear compartmentalization in yeast has been mostly studied in nutrient-replete conditions; however, yeast in the wild can experience more challenging growth conditions and have developed ways to cope with the absence of nutrients. In a liquid culture, yeast cells undergo multiple metabolic transitions from exponential phase to stationary phase (SP). During exponential growth in rich medium containing glucose as a carbon source (Yeast Peptone Dextrose, YPD), yeast cells are fermenting glucose, their preferred source of energy. When glucose is exhausted, the cells enter the diauxic shift (DS), a phase in which they are switching from fermentation to respiration using fermentation products such as ethanol as a carbon source, drastically slowing down their growth rate. When ethanol is exhausted, cells will stop dividing and enter SP. Two types of cells are found in SP cultures and are separable by density gradient - low density (LD) and high-density (HD) populations. The LD population is composed mainly by senescent and apoptotic cells (non-quiescent cells, NQ), while the high-density (HD) fraction is constituted essentially by quiescent cells (Q) (Allen *et al*, 2006). Q and NQ cells have different properties: NQ cells are short-lived, asynchronously arrested and replicatively old cells, while Q cells are long-lived, G_0_-arrested and unbudded “virgin” cells. Q cells are more resistant to various stress and maintain the property to restart the cell cycle after long time periods after addition of the missing nutrients (Gray *et al*, 2004; De Virgilio, 2012; Sun & Gresham, 2021; Breeden & Tsukiyama, 2022). Q cells undergo drastic spatial reorganization of their components when they transition from exponential phase to stationary phase, both in the cytoplasm (proteasome, actin cables, microtubules, mitochondria) and in the nucleus (Sagot & Laporte, 2019; Breeden & Tsukiyama, 2022). The genome is drastically rearranged: the nuclear and nucleolar volume is reduced (Johnston *et al*, 1977; Guidi *et al*, 2015; Wang *et al*, 2016); chromatin is compacted (Schafer *et al*, 2008); promoters are deacetylated and transcription is shut down (Mews *et al*, 2014; McKnight *et al*, 2015; Young *et al*, 2017); and RNAPII relocalizes to a subset of intergenic promoters (Radonjic *et al*, 2005; Baquero Pérez *et al*, 2025).

We have previously shown that telomere spatial organization is drastically affected by metabolic transitions in a reversible manner. While telomeres are organized into 3 to 5 foci at the nuclear periphery in exponential phase, they become brighter and fewer in number a few hours after the diauxic shift when cells have switched their metabolism from fermentation to respiration. Upon entry into stationary phase, telomere foci group together into a hypercluster (HC) in the center of the nucleus in Q cells while their spatial organization is not evolving in non-quiescent cells, making HC a landmark of quiescent cells (Guidi *et al*, 2015). In addition, we showed that HC contributes to the survival of Q cells (Guidi *et al*, 2015). This reorganization of telomeres is rapidly reversed upon release into rich medium indicating that the level of telomere clustering is tightly regulated and coordinated with carbon source availability (Baquero Pérez *et al*, 2025). We showed that the Sir3 protein is required for the formation of the telomere HC in quiescence (Guidi *et al*, 2015); however the mechanisms regulating the clustering of telomeres upon metabolic transitions have yet to be identified.

Here we investigate the mechanisms regulating the spatial organization of telomeres during entry into quiescence by combining genetics and physical modeling. We show that Sir3 is subjected to post-translational modifications (PTMs) during these metabolic transitions; however, these PTMs have a modest effect on telomere clustering. Our physical modeling reveals that telomere clustering and telomere anchoring can have antagonistic effects and predicts that decreasing telomere anchoring is more effective to drive the formation of HC than increasing interactions between telomeres. In good agreement with our model, deletion of *ESC1* leads to the precocious appearance of telomere hyperclusters after the diauxic shift and in NQ cells in stationary phase. Furthermore, Esc1 ability to maintain telomeres at the nuclear periphery in early post-diauxic shift cells is abolished by mutating serine 1450 of Esc1, whose phosphorylation is required *in vitro* for its interaction with the HBRCT domain of Sir4 (Deshpande *et al*, 2019). Modifying this unique residue phenocopies deletion of *ESC1* in post-diauxic shift cells, strongly indicating that Esc1 mediated anchoring is regulated by phosphorylation upon metabolic transition, driving massive change in telomere clustering.

## Results

### Sir3 levels and post-translational modifications are not required for the formation of telomere hyperclusters during stationary phase

Telomere clustering is highly dynamic and regulated upon metabolic transitions (Guidi *et al*, 2015; Laporte *et al*, 2016), however the molecular mechanisms driving the spatial reorganization of telomeres upon glucose exhaustion have not yet been identified. An obvious candidate for the regulation of telomere clustering during metabolic transitions is a member of the Sir complex, Sir3. Indeed, we showed that Sir3 is actively involved in the formation of telomere foci through Sir3-Sir3 interactions (Ruault *et al*, 2011, 2021). Sir3 overexpression leads to the formation of telomere hyperclusters which strikingly resemble those observed in quiescence (**Figure 1A**). However, Sir3 protein levels are stable in stationary phase (Guidi *et al*, 2015, **Figure 1B**) ruling out the simple hypothesis that telomere hyperclustering in quiescent cells is due to a strong increase in Sir3 levels. Laporte *et al*. (2016) observed that the BY strain background is less efficient than W303 to form telomere hyperclusters in stationary phase and suggested that this difference could be due to a ∼1.5-fold increase in Sir3 levels in W303 compared to BY. To test this hypothesis, we introduced additional *SIR3* copies in the genome and quantified the clustering in the modified strains using an image analysis pipeline developed in house (NuFoQal, see Methods). We confirmed that the formation of telomere hyperclusters is less frequent in the BY strain than in W303 during stationary phase (68% in W303 vs. 20% in BY; χ^2^, p <0.0001; **Figure S1A**). However, increasing Sir3 levels up to 3.5-fold did not rescue this defect (**Figure 1C, Figure S1B**). Therefore, we conclude that Sir3 protein levels is not responsible for the scarcity of telomere hyperclusters in the BY cells; rather, the difference may result from their defects to enter quiescence (Gaisne *et al*, 1999; Miles *et al*, 2023).

**Figure 1:**
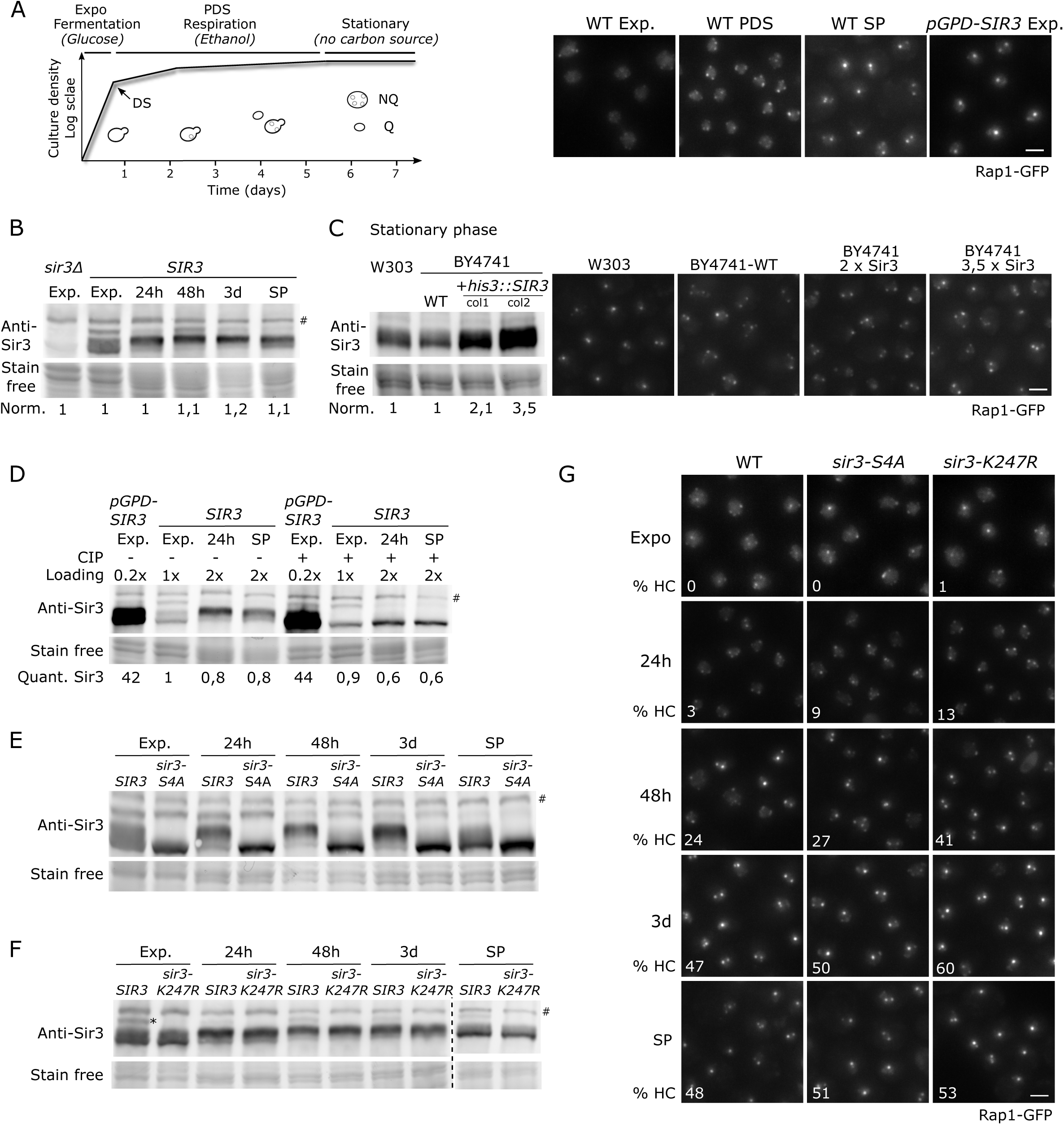
Sir3 levels and modifications do not impact the formation of telomere hyperclusters during stationary phase. A) Left: Growth curve of *S. cerevisiae* grown in liquid glucose-based rich medium. Cells are first growing exponentially, fermenting glucose and producing ethanol. After glucose exhaustion (Diauxic Shift, DS), cells use ethanol as a carbon source (Respiration). When all carbon sources are exhausted, cells enter stationary phase. At this stage, cultures are heterogenous, composed of quiescent cells (Q cells) and non-quiescent cells (NQ cells). Right: Representative fluorescent images of the telomere-associated protein Rap1 tagged with GFP (yAT2487) during exponential phase, after the post-diauxic-shift (PDS), during stationary and in exponentially growing cells overexpressing Sir3 (yAT3110). Scale bar is 2µm. Western blots in B, C, D, E, F were done using a Sir3 antibody. The # labels an aspecific band detected by the anti-Sir3 antibody. The total amount of Sir3 protein per lane was normalized to the stain-free signal per lane. B) Sir3 levels were assayed in a *sir3Δ* (yAT2553) and in the WT strain (yAT2487) during exponential phase and at 24h, 48h, 3 days and 7 days for the WT. C) WT W303 (yAT2487), WT BY4741 (yAT1950) and WT BY4741 transformed with additional *SIR3* copies (col1: yAT4773; col2: yAT4818) were grown to stationary phase. Left: Immunobloting using the Sir3 antibody; Right: Representative fluorescent images of the strains. Scale bar is 2µm. D) Sir3 was detected by immunoblotting in crude extracts, with or without CIP (phosphatase) treatment, in the Sir3-overexpressing strain (yAT3110) during exponential phase, as well as in the WT strain (yAT2487) during exponential phase, at 24h and 7days. E) Sir3 was detected in crude extracts from WT (yAT2487) and from the *sir3-S4A* mutant (yAT3905) during exponential phase, at 24h, 48h, 3 days and 7days. F) Same as in E) with crude extracts from WT (yAT2487) and from the *sir3-K247R* mutant (yAT3868). The asterisk indicates the sumoylated-Sir3 protein. G) Representative images of Rap1-GFP samples from WT (yAT2487), *sir3-S4A* mutant (yAT3905) and *sir3-K247R* mutant (yAT3868). White numbers at the bottom left of each images represent the percentage of cells with a hypercluster. Scale bar is 2µm.

While Sir3 levels are globally stable during metabolic transitions, we observed some variations in Sir3 migration patterns that reflect post-translational modifications. Indeed, three main Sir3 species are observed on western blots from exponential phase cells: a fast, an intermediary and a slow migrating band (**Figure 1B, D, E, F**). After 24h of culture and at later time points, the highest and lowest Sir3 migrating bands decrease relative to the intermediary band (**Figure 1B, D, E, F**). It was previously shown that Sir3 is phosphorylated (Stone & Pillus, 1996; Ai *et al*, 2002; **Figure S1C**). To determine whether one of these bands was due to Sir3 hyperphosphorylation, we digested the total protein extracts with the calf intestinal alkaline phosphatase (CIP). The phosphatase treatment caused the disappearance of the intermediate Sir3 band to the benefit of the fastest migrating band (**Figure 1D**). Thus, Sir3 is hyperphosphorylated when cells are switched from fermentation to respiration, the highest levels of Sir3 phosphorylation being reached between 48h and 3 days and decreasing slightly at later time points (**Figure 1B**). To determine whether the phosphorylation of Sir3 during metabolic transitions was involved in the formation of telomere hyperclusters, we assessed telomere clustering in the *sir3-S4A* mutant described by Ray *et al*, 2003 as a non-phosphorylatable form of Sir3. Sir3-S275A-S282A-S289A-S295Ap exhibits a migration profile similar to the dephosphorylated wild-type protein (**Figure 1E, Figure S1D**) but has no strong negative effect on the kinetics of telomere clustering (**Figure 1G**). On the contrary, we observed a small positive effect of the *sir3-S4A* mutation on telomere clustering at the beginning of glucose deprivation (3% of HC in the WT at 24h vs 9% in the *sir3-S4A* mutant) indicating that the phosphorylation is rather counteracting clustering **(Figure S1E** for quantifications). Sir3 is phosphorylated on other serine/threonine residues (our unpublished data and GPMdb : Generalized Proteomics data Meta-analysis database - Craig *et al*, 2004). Therefore, we constructed a *sir3* allele in which we mutated 9 other serine/threonine frequently phosphorylated in Sir3 (S315A-S322A-S323A-S327A-T335A- S336A-S339A-S340A-S344A). The Sir3 migration profile of this mutant, *sir3-S8A-T1A*, is identical to WT Sir3 and the dynamics of telomere clustering in this strain is not affected compared to the wild-type strain (**Figure S1F**).

We then asked whether the modification corresponding to the slowest-migrating Sir3 band was important for telomere clustering. We hypothesized that this band could result from Sir3 sumoylation since Sir3 was reported to be sumoylated on lysine 247 (R. Sternglanz, personal communication). The slowest Sir3 migrating band disappeared when the lysine 247 was mutated to arginine, indicating that Sir3 is likely sumoylated at this amino acid (**Figure 1F**). Sir3 sumoylation is mainly observed at the beginning of the kinetics, always present in exponential phase and decreasing between 24h and 48h (**Figure 1D, E, F**). Blocking Sir3 sumoylation did not prevent the formation of telomere hyperclusters in quiescent cells (**Figure 1G**), on the contrary, we observed an increase in the percentage of cells with a hypercluster from 24h to 3 days in this mutant indicating that Sir3 sumoylation is rather a negative regulator of telomere hypercluster formation during metabolic transitions (**Figure 1G, Figure S1E** for quantifications). Thus, Sir3 post-translational modifications, both phosphorylation and sumoylation are not required for telomere hypercluster formation; rather they have a modest negative impact.

### Physical modeling indicates that telomere anchoring and telomere clustering antagonize each other

To better understand the physical constraints affecting the phenomenology and regulation of telomere clusters in the nucleus, we adapted a previously developed physical model of chromosomes in the yeast nucleus. *S. cerevisiae* was the first eukaryote whose general nuclear architectural determinants were reproduced by coarse-grained Molecular Dynamics simulations (Wong *et al*, 2012; Hajjoul *et al*, 2013; Arbona *et al*, 2017). These simulations modelled the yeast chromosomes as bead-spring polymers, confined in a hollow sphere representing the cell nucleus, with specific interactions at centromeres mimicking the effects of microtubules and maintaining them in proximity of the spindle pole body. Further, the ribosomal DNA was modeled by a chain of larger beads inserted on the right arm of chromosome XII, that, due to their different size and the geometry of confinement, naturally segregate on the opposite side of the spindle pole body. In those models, the telomeres are permanently associated to the nuclear envelope (**Figure 2A**). However, *in vivo,* telomeres are dynamic and not always associated to the nuclear periphery (Tham *et al*, 2001; Hediger *et al*, 2002). Furthermore, random collisions between telomeres alone cannot account for the Sir3-dependent variability observed in cluster formation (Hozé *et al*, 2013) or for the emergence of telomere hyperclusters observed in quiescent cells (Guidi *et al*, 2015). To account for those observations, we modified the initial model by introducing: i) attractive telomere-to-telomere interactions proportional to *∈*_*TT*_, representing the glueing effect of Sir3 (Ruault *et al*, 2011, 2021) ; ii) variable telomere-to-nuclear envelope interactions proportional to *∈*_*TE*_, representing the effects of anchoring of telomeres to the nuclear envelope iii) and, finally, rDNA-to-nuclear envelope fixed interactions meant to prevent telomeres to locate in the region of the nuclear envelope associated to the nucleolus (Therizols *et al*, 2010). The new model is simulated at a resolution of 1 bead per ∼ 800 base pairs. The two variables *∈*_*TT*_ and *∈*_*TE*_ respectively tune the clustering between telomeres and their anchoring to the nuclear envelope. We used this new model to test the relationship between telomere anchoring and clustering.

**Figure 2:**
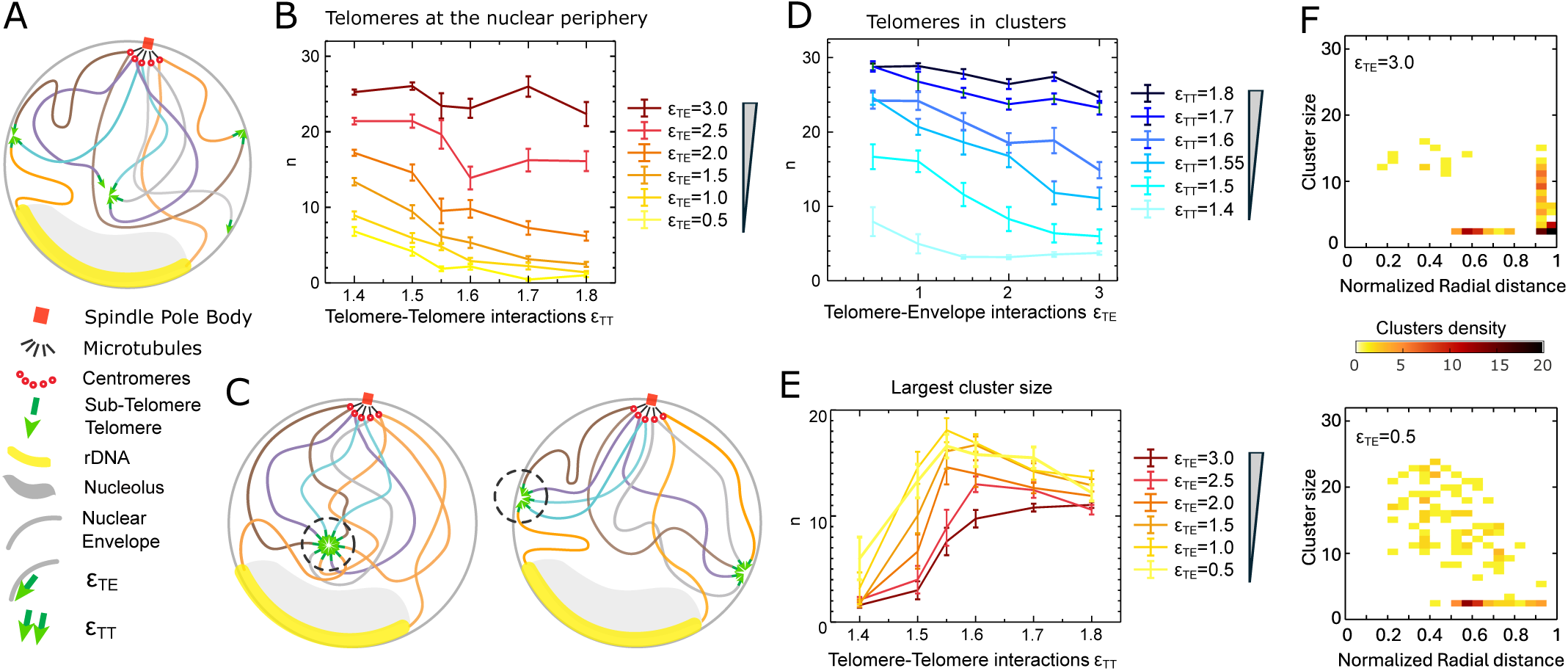
Molecular dynamics simulations suggests that dissociation from the nuclear periphery is sufficient to drive the formation of telomere hyperclusters. All plots and legends are done in function of either telomere-telomere attractive interactions, or telomere-nuclear envelope attractive interactions. A) Main structural elements implemented in the model, namely the spindle pole body, the chromosomes, the nucleolus and the telomeres. B) Number of telomeres at the nuclear periphery as a function of telomere-telomere attractive interactions. C) Schematic representation of the steric hindrance of chromosome arms emerging from telomere clusters forming a polymer brush (dotted line circle) extending from a collapsed telomeric cluster. The volume available to the brush is reduced when it is anchored to the nuclear envelope. D) Number of telomeres belonging to a cluster of size greater or equal to 2, as a function of telomere-nuclear envelope attractive interactions. E) Size of the largest cluster as a function of telomere-telomere interaction. F) Heat-map of all clusters identified in the simulations (60 replicas) indicating their radial distance respect to the center of the nucleus and their size.

All simulations started from a conformation sampled from a condition where both *∈*_*TT*_ and *∈*_*TE*_ are set to zero. The coarsening of telomere clusters and their positioning at the nuclear envelope is observed over a timescale compatible to a full cell cycle. Our results show that the positioning of telomeres at the nuclear periphery mainly depends on *∈*_*TE*_, as expected. However, for values of *∈*_*TE*_ lower then ≈ 3k_B_T, it also depends on *∈*_*TT*_: with telomeres detaching more from the nuclear envelope for higher levels of clustering as *∈*_*TT*_ increases (**Figure 2B**). We attribute this effect to the steric repulsion of chromosome arms emerging from telomere clusters forming the so called polymer brush, as observed on Hi-C contact maps for centromeres (Daoud & Cotton, 1982; Wong et al, 2012; Goloborodko et al, 2013; Arbona et al, 2017) . The space being more limited at the nuclear border that in the center of the nucleus, this prevents the formation of large clusters **(Figure 2C and 2D)**. Reciprocally, the number of telomeres in clusters mainly depends on *∈*_*TT*_, but for values of *∈*_*TT*_ lower than ≈ 1.7k_B_T, increasing *∈*_*TE*_ decreases the proportion of telomeres within clusters **(Figure 2D).** In other words, strong telomere anchoring tends to fragment weak clusters.

Focusing on the size of the biggest cluster, we observe that the highest value is reached for intermediate values of *∈*_*TT*_, specifically around *∈*_*TT*_ ≈ 1.55k_B_T. Although counterintuitive, this can be explained, again, by the repulsion between chromosome arms preventing stable clusters to merge into a hypercluster. In contrast, for intermediate values of *∈*_*TT*_, individual telomeres can detach from small clusters and join bigger clusters that are more stable, leading to the formation of stable hyperclusters **(Figure 2E)** (Bunin & Kardar, 2015; Scolari *et al*, 2018). This phenomenon is analogous to Ostwald ripening in phase separation (Ostwald, 1897; Zwicker *et al*, 2025).

We conclude that *in vivo*, values of *∈*_*TT*_should be near critical (≈ 1.55k_B_T) to allow hypercluster formation. Taking this value for *∈*_*TT*_, we compared cluster sizes and their position for high or low value of *∈*_*TE*_(*∈*_*TE*_ = 3.0*k*_*B*_*T vs ∈*_*TE*_ = 0.5*k*_*B*_*T*). We observe that in the latter, large clusters form only far from the nuclear envelope while there are no cluster containing more than half of telomeres in the former (**Figure 2F**). Therefore, in our model, dissociation from the nuclear periphery is sufficient to drive the formation of telomere hyperclusters without changing the strength of telomere-telomere interactions, provided that these interactions are not too strong **(Figure S2)**.

### Esc1 telomere anchoring is essential to prevent telomere hyperclustering after the diauxic shift

Our physical modeling prompted us to test the impact of telomere anchors on telomere clustering. As mentioned above telomeres are anchored to the nuclear periphery by multiple redundant pathways (Taddei *et al*, 2004; Bupp *et al*, 2007; Schober *et al*, 2009; Chan *et al*, 2011; Park *et al*, 2011; Van de Vosse *et al*, 2013; Lapetina *et al*, 2017). We thus inactivated or deleted some of these anchors or their regulators and imaged the Rap1-GFP telomeric protein in those mutants after metabolic transitions (during exponential phase, a few hours after the diauxic shift and later at 24h). Most of the mutations / deletions did not impact telomere clustering during the time course (*heh1Δ heh2Δ*, *mps3Δ75-150*, *mlp1Δ mlp2Δ*, *yku70Δ yku80Δ rif1Δ rif2Δ* or *pfa4Δ*) or had a negative effect on telomere clustering (*nup170Δ)* **(Figure 3A)**. In contrast, *ESC1* deletion had a positive impact on telomere clustering **(Figure 3A).** Indeed, in an *esc1Δ* strain, telomere hyperclusters are detected earlier and are more frequent than in wild type after the diauxic shift (**Figure 3B**). A few hours after the diauxic shift 5% of the *esc1Δ* cells harbor hyperclusters while only 1.6% of the WT cells have such a structure (χ^2^, p = 0.0001). At 24h, the differences are more pronounced, with 53% of the *esc1Δ* cells with at least one hypercluster, but only 9% for the WT (χ^2^, p <0.0001). In addition, the proportion of cells with only 1 or 2 clusters is strongly increased in the *esc1Δ* at 24h (73% for *esc1Δ* versus 25% for the WT - pie charts from **Figure 3B**; χ^2^, p <0.0001). Fluorescent In Situ Hybridization (FISH) experiments using a Y’ labelled probe confirmed that the bright clusters indeed result from the clustering of telomeres **(Figure S3A)**. By measuring the distance between the nuclear envelope and telomere foci, we showed that telomere clusters of the *esc1Δ* strain are less peripheral than the ones of the WT (**Graph Figure 3C**). Furthermore, we observed that the brightest clusters of *esc1Δ* cells localize in the center of the nucleus, like hyperclusters in quiescent cells (**images Figure 3C)**. Since telomere hyperclusters appear earlier in the *esc1Δ* compared to the wild-type during metabolic transitions, we hypothesized that *esc1Δ* strains may have an accelerated metabolism (exhausting glucose earlier than wild-type). However, the kinetics of endocytosis of Hxt1p (a glucose transporter expressed when glucose is high) and the expression of Hxt5p (a glucose transporter expressed when glucose is low) in *esc1Δ* strains is like wild type, ruling out this hypothesis (**Figure S3B**). Next, since telomeres have greater chance to encounter and interact in a smaller volume, we investigated the potential role of nuclear size on the formation of telomere hyperclusters. The sizes of the nuclei are very similar in WT and *esc1Δ* cells in exponential phase (2µm). At 24h, the nucleus size of unbudded cells with a diameter below 4.5µm shrank significantly to 1.6 µm both in WT and *esc1Δ* cells while the size of the nucleus did not change in larger cells (diameter of cells >4.5µm). Thus, precocious telomere hyperclustering of the *esc1Δ* strain is not due to a decreased nuclear volume of *esc1Δ* cells (**Figure S3C, S3D**).

**Figure 3:**
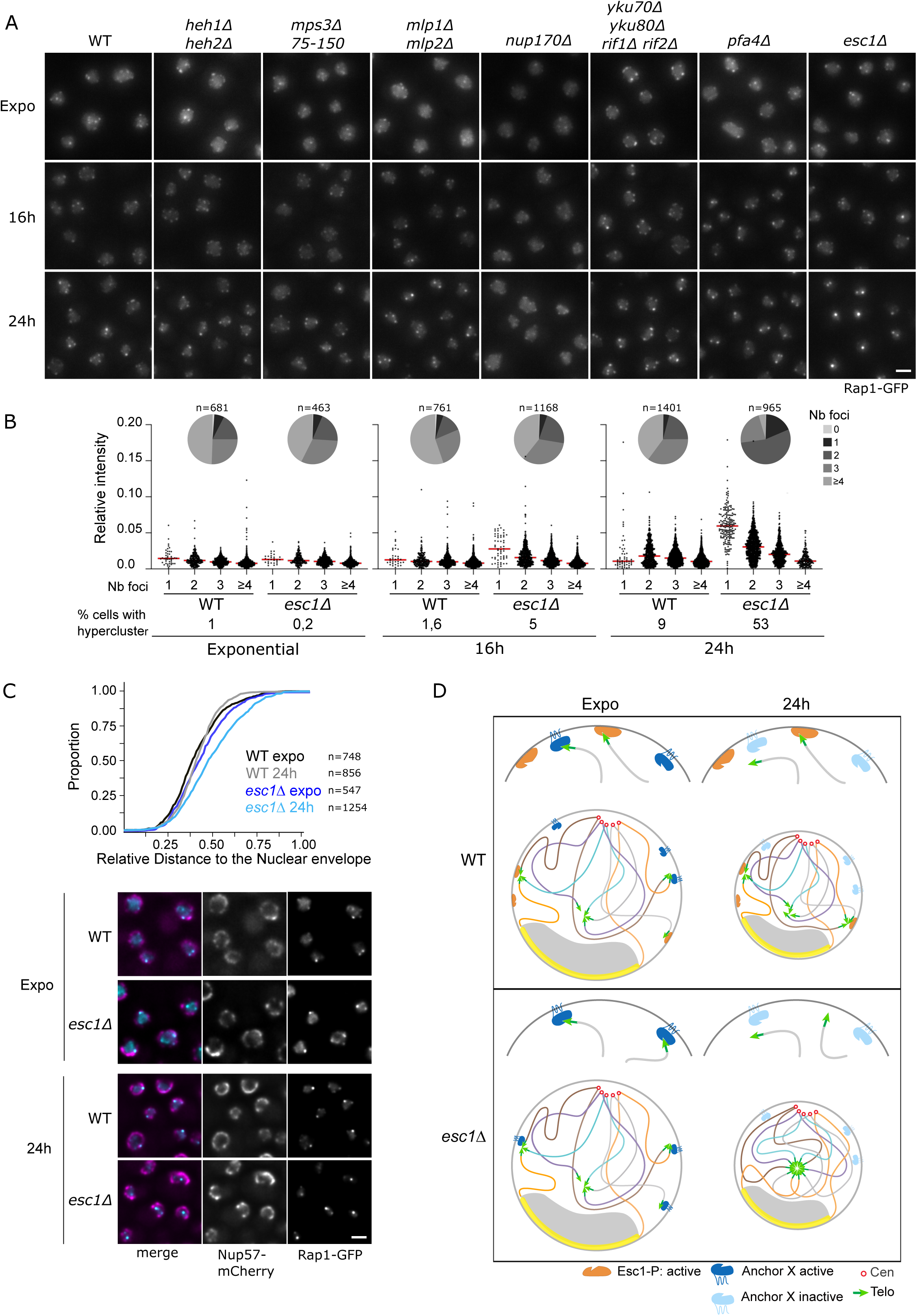
Esc1 is the main active anchor a few hours after glucose exhaustion and prevent the formation of telomere hyperclusters in the center of the nucleus. A) Representative fluorescent images of Rap1-GFP strains during exponential phase, at 16h and 24h: WT (yAT2487), *heh1Δ heh2Δ* (yAT3814), *mps3Δ75-150* (yAT3895), *mlp1Δ mlp2Δ* (yAT4432), *nup170Δ* (yAT4014), *yku70Δ yku80Δ rif1Δ rif2Δ* (yAT4396), *pfa4Δ* (yAT4903) and *esc1Δ* (yAT3811). Scale bar is 2µm. B) Quantifications of telomere clustering: Dot plots represent the cluster intensity relative to the total nuclear signal in cells with 1, 2, 3 or 4 and more foci (red bar is the median). The percentage of hyperclusters is indicated for each strain at the bottom of the graph. The pie charts represent the distribution of the cells according to the number of foci per cell, n is the total number of cells analyzed. C) Cumulative distributions of the distance Telomere cluster – Nuclear periphery in WT (yAT4606) and *esc1Δ* (yAT4607) during exponential phase and at 24h. Representative deconvolved fluorescent images of the cultures used to generate the graph. Scale bar is 2µm. D) Schematic representation of a nucleus during exponential phase and at 24h in WT and *esc1Δ* cells.

We thus conclude that Esc1 is the main telomere anchor after the diauxic shift. In exponential phase, in rapid cycling cells, the existence of one or multiple redundant anchoring pathways prevents the release of telomeres from the nuclear envelope and their clustering in the nuclear center in the absence of Esc1. We hypothesize that this or these pathways (that we named anchor X) are impaired after the diauxic shift, leaving Esc1 as the main available anchor to prevent hyperclustering of telomeres at 24h (**Figure 3D**).

### Esc1-independent anchoring is regulated by glucose signaling

Telomere hyperclusters are not only forming in Q cells that have reached stationary phase (glucose exhaustion), but they can also be observed in cells starved for 24h after the diauxic shift (PDS starvation). On the contrary, starvation of exponentially growing cells does not cause the formation of telomere hyperclusters after 24h of starvation (Guidi *et al*, 2015). We asked whether anchoring through Esc1 could be the mechanism that prevents the formation of telomere hyperclusters in cells starved from exponential phase. Strikingly, the *esc1Δ* strain formed bright telomere hyperclusters between 7h and 20h of starvation contrary to the WT (**Figure 4A**). Thus, when exponentially growing cells are abruptly starved in water, the Esc1-independent anchoring pathway is altered and, in some cells, Esc1 is the main anchor left.

**Figure 4:**
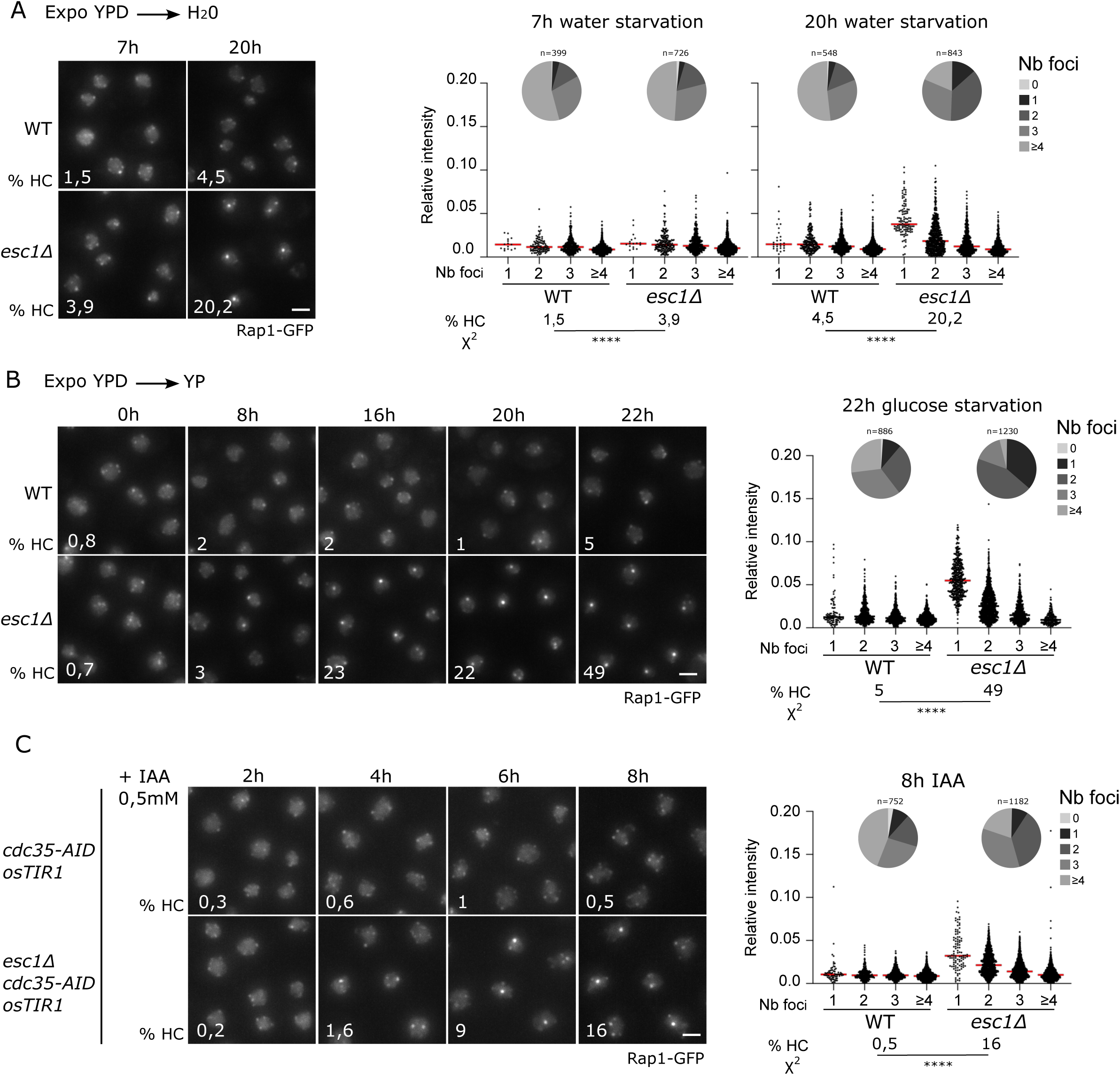
Esc1-independent anchoring is inactivated in the absence of glucose. A) B) C) Representative fluorescent images of the telomere-associated protein Rap1 tagged with GFP in WT and *esc1Δ.* Quantifications of telomere clustering: Dot plots represent the cluster intensity relative to the total nuclear signal in cells with 1, 2, 3 or 4 and more foci (red bar is the median). The percentage of cells with hyperclusters is indicated for each strain on the corresponding fluorescent image. Statistical analysis was done using a two-sided Chi-square test with a confidence interval of 99%; **** = p < 0.0001. The pie charts represent the distribution of the cells according to the number of foci per cell, n is the total number of cells analyzed. A) Cells were grown to exponential phase in YPD (OD_600nm_ =1) and switched to water. Images were taken at 7h and 20h. B) Cells were grown to exponential phase in YPD (OD_600nm_ =1) and switched to YP medium (YPD deprived of glucose). Images were taken every 2h from 0h to 22h. C) Strains expressing Rap1-GFP, Cdc35-AID, TIR1p under the control of the GPD promoter with or without the *ESC1* gene (yAT3715 and yAT4580 respectively) were grown to OD_600nm_ = 0,4. 0,5 mM IAA was added to the cultures at time 0h. Images were taken every 2h from 2 to 8h of culture.

As glucose plays a central role in the regulation of telomere anchoring, we decided to test directly the impact of glucose starvation on the formation of telomere hyperclusters by switching exponentially growing cells to YP (Yeast Peptone medium without glucose) and monitor the clustering of telomeres for 22h. In the WT strain, the starvation of exponentially growing cells from glucose has no impact on telomere clustering. However, in the *esc1Δ* strain, telomere clusters begin to group in large clusters after 8h of glucose starvation. In the *esc1Δ* strain, the percentage of cells with a hypercluster increases with the duration of glucose deprivation, reaching 49% at 22h (only 5% in the WT) (**Figure 4B**, detailed quantifications **Figure S4A**). Thus, we can conclude that the Esc1-independent anchoring mechanism, which is active in exponentially growing cells is progressively lost upon glucose starvation. Next, we asked whether this was due to the absence of glucose or to the change of carbon source by switching the cells from YPD to YP-EtOH (YP 2% Ethanol). Under these conditions, *esc1Δ* cells are forming telomere hyperclusters similar to those observed in YP or in water (**Figure S4B**). This result indicates that the Esc1-independent anchoring mechanism is deactivated specifically in the absence of glucose.

Glucose signaling involves multiple pathways that adjust the growth rate of a culture based on glucose availability. The Ras/PKA pathway plays a key role in coordinating the cellular response to glucose levels. When glucose is present, Ras activates the Cdc35 (Cyr1) adenylate cyclase leading to the production of cAMP and the subsequent activation of PKA (Zaman *et al*, 2009; Peeters & Thevelein, 2014). To test whether Esc1-independent anchoring is regulated by the Ras/PKA pathway, we inactivated the adenylate cyclase using a Cdc35-degron (Waterman et al., in preparation). In cells expressing Rap1-GFP, Cdc35 fused to the AID tag and *TIR1* under the control of a strong promoter (*GPD-TIR1*) were grown to exponential phase and auxin was added to the culture, images were taken every two hours after auxin addition. As expected, both wild-type and *esc1Δ* cells accumulate in G1 phase (Matsumoto *et al*, 1983; Werner-Washburne & Singer, 1993). However, cells grow significantly during the eight hours after Cdc35 depletion, contrary to the cells starved in water or YP, indicating that these cells are still metabolically active (**Figure S4C**). Yet, cells did not exhaust the glucose during this time (**Figure S4D**). Contrary to the wild-type cells, after 6h of Cdc35 inactivation, *esc1Δ* cells form telomere hyperclusters (9% in *esc1Δ* vs 1% in WT; χ^2^, p <0.0001) and this percentage was increased after 8h (16% in *esc1Δ* vs 0,5% in WT; χ^2^, p <0.0001) (**Figure 4C**, detailed quantifications **Figure S4E**). We obtained similar results with Sir4-pH-tdGFP tagged strains upon Cdc35 inactivation (**Figure S4F**). Thus, the Esc1-independent anchoring pathway is inactived when the Ras/PKA pathway is inhibited. Because PKA-inactivated cells also arrest in G1 (Werner-Washburne & Singer, 1993), we examined whether a G1-arrest induced by alpha-factor could lead to the formation of telomere hyperclusters in the absence of Esc1. G1-arrested *esc1Δ* cells formed brighter clusters than WT, although these clusters were not as bright as those observed upon Cdc35 inactivation **(Figure S4G)**. The percentage of cells with a hypercluster was significantly increased in *esc1Δ* cells treated for 6h with alpha-factor compared to WT (4,4% in *esc1Δ* vs 1,4% in WT; χ^2^, p = 0.0253). Prolonged alpha-factor arrest leads to the deformation of nuclei complicating the interpretation (light-transmitted images **Figure S4G**). However, our data suggest that G1 arrest alone is not sufficient to inactivate Esc1-independent anchoring, indicating that inhibition of the PKA pathway contributes independently to this process.

The Esc1-independent anchoring pathway is thus tightly regulated by metabolism and cell cycle in exponential phase. To identify the redundant anchors at play in exponential phase together with Esc1, we created multiple strains in which Esc1, and some potential anchors or regulators of the anchors were deleted; however, we did not observe telomere hyperclusters in exponential phase in any of these strains (**Table 1**). In conclusion, telomere anchoring at the nuclear periphery is ensured by at least two redundant pathways in exponential phase; i) Esc1 and ii) one or several unidentified pathways. Those redundant anchors are inactivated when cells are deprived of glucose or upon Cdc35 depletion, while Esc1 remains active in those conditions.

**Table 1:**
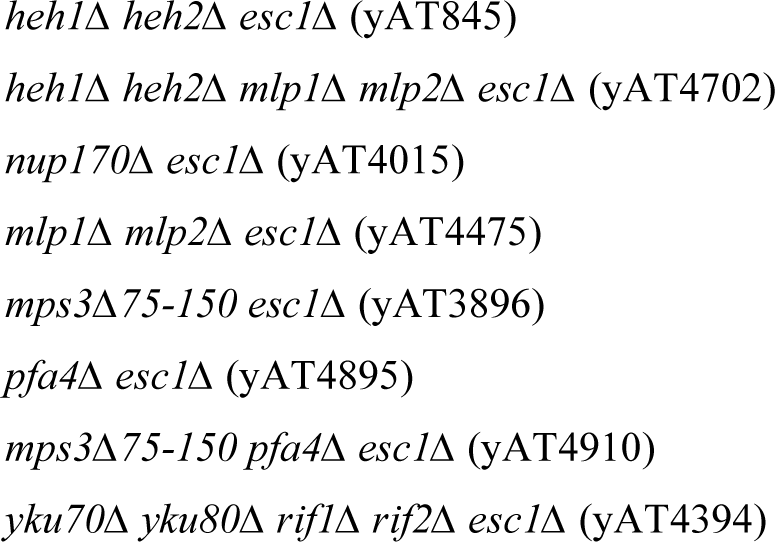
List of multiple anchoring mutants tested for telomere hyperclustering in exponential phase.

### Esc1 is the anchor that prevents telomere clustering in non-quiescent cells

In cultures undergoing carbon source exhaustion, Esc1 is the main telomeric anchor at play a few hours after the diauxic shift and is efficiently counteracting telomere clustering at this stage in most of the cells. Since telomere hyperclusters form specifically in wild-type Q cells when they reach SP, we reasoned that the Esc1 anchoring pathway must be inactivated specifically in Q cells to allow the formation of telomere hyperclusters. Monitoring telomere foci from the exponential phase to stationary phase (6-7days) revealed that telomere hyperclusters accumulation in *esc1Δ* cells occurs faster than in WT cells and in a higher proportion of cells (86% vs 52%; χ^2^, p <0.0001) (**Figure 5A**). In addition, the proportion of cells with only one focus was higher in the *esc1Δ* (61%) compared to the WT (34%) (pie charts) and the brightness of their clusters was also significantly higher in the *esc1Δ* compared to the WT (median at 7,8% of the total Rap1-GFP signal in *esc1Δ* vs 5,9% in the WT; Welch’s t test, p < 0.0001). We observed similar results by monitoring three other telomere bound proteins during metabolic transitions: Yku80, Sir3 and Sir4 **(Figure S5A)**. Deletion of *ESC1* did not cause any variations in Sir3 levels or post-translational modifications compared to WT during the entire kinetics **(Figure S5B)** and did not change the size of the nucleus **(Figure S5C)**.

**Figure 5:**
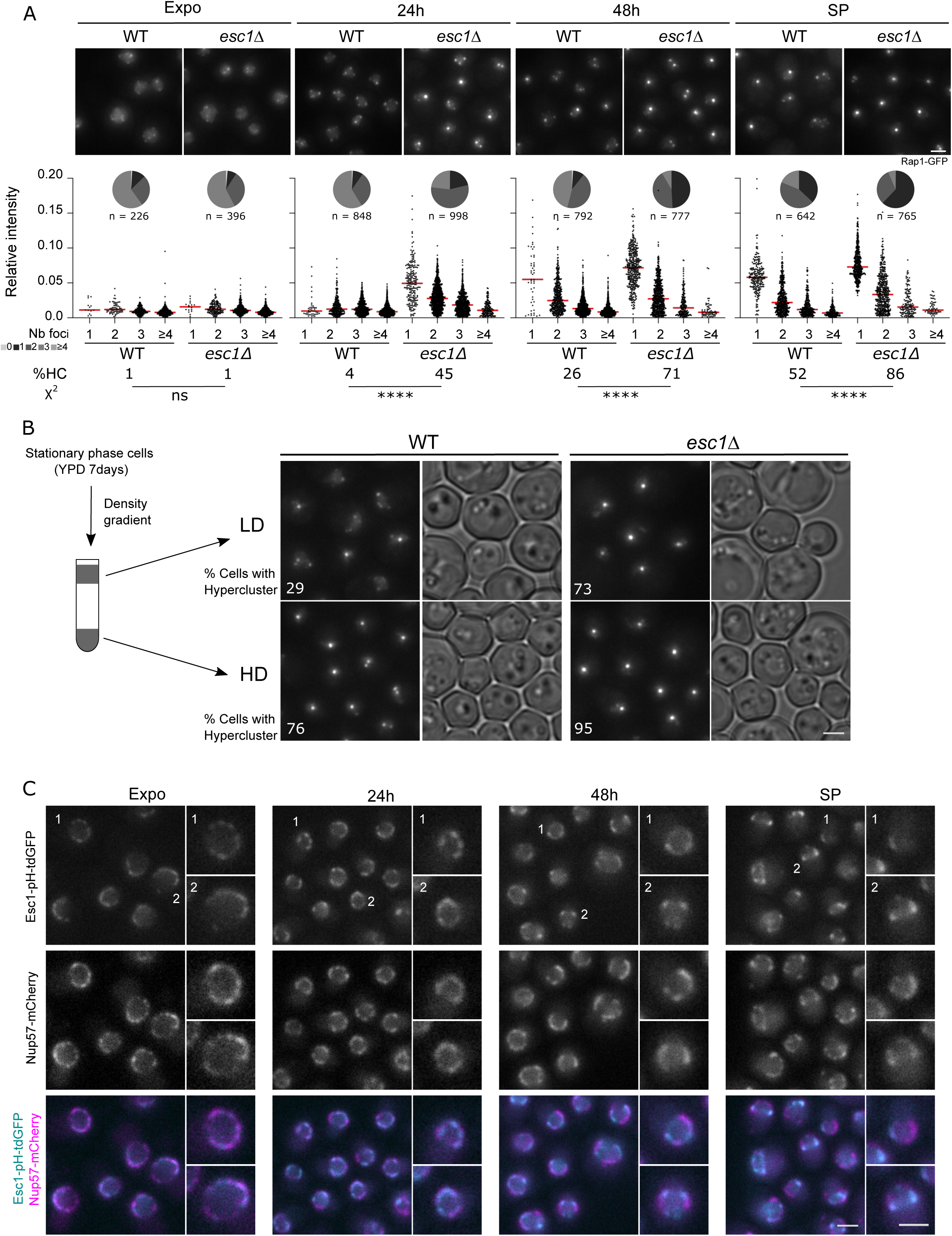
Esc1 prevents the formation of telomere hyperclusters in non-quiescent cells. A) Representative fluorescent images of Rap1-GFP WT (yAT2487) and *esc1Δ* (yAT3811) strains during exponential phase, at 24h, 48h and during stationary phase. Scale bar is 2µm. Quantifications of telomere clustering: Dot plots represent the cluster intensity relative to the total nuclear signal in cells with 1, 2, 3 or 4 and more foci (red bar is the median). The percentage of cells with a hypercluster is indicated at the bottom of each image. Statistical analysis was done using a two-sided Chi-square test with a confidence interval of 99%; **** = p < 0.0001. The pie charts represent the distribution of the cells according to the number of foci per cell, n is the total number of cells analyzed. B) Schematic representation of a density gradient. WT (yAT2487) and *esc1Δ* (yAT3811) strains expressing Rap1-GFP were grown to stationary phase and were loaded on a density gradient. Representative fluorescent and light transmitted images of the cells recovered from the LD (low-dense) and the HD (High dense) fractions of the gradients. Scale bar is 2µm. C) Representative fluorescent images of a strain expressing Esc1-pHtdGFP and Nup57-mCherry (yAT4771) at different time points of the culture: exponential phase, 24h, 48h and stationary phase. Scale bar is 2µm.

We next asked whether the highest proportion of cells with hyperclusters in the *esc1Δ* was due to an increased rate of forming Q cells in *esc1Δ,* by sorting HD vs LD fraction of *esc1Δ* and WT stationary phase cultures on density gradients. The fraction of HD cells is similar in WT and *esc1Δ,* excluding the hypothesis that *esc1Δ* cells reach the quiescent state more efficiently than WT (data not shown). However, a large fraction of the LD (enriched in NQ) *esc1Δ* cells harbor telomere hyperclusters (73% of the cells with a HC in *esc1Δ* versus 29% in a WT, χ^2^, p <0.0001) **(Figure 5B, Figure S5D),** explaining the high percentage of cells with a hypercluster in the *esc1Δ* unsorted stationary phase cultures. Thus, the loss of the anchoring in *esc1Δ* NQ cells leads to the formation of telomere hyperclusters in the center of the nucleus, similarly to what we observed in wild-type Q cells. Therefore, in wild-type, Esc1 dependent telomere anchoring is maintained in NQ cells and lost specifically in Q cells.

We next tested whether Esc1 levels and/or localization are modified during metabolic transitions and differ between Q and NQ cells by tagging Esc1 with a fluorescent tag (pH-tdGFP) in a strain expressing Nup57 tagged with mCherry to label the nuclear pores. Imaging of Esc1-pH-tdGFP shows that Esc1 levels are very similar during the whole time-course and that Esc1 remain localized at the nuclear periphery at any timepoint (**Figure 5C**). However, we did notice subtle changes of Esc1 localization during metabolic transitions; specifically, Esc1 was evenly distributed along the nuclear membrane, excluding the nucleolar region, during logarithmic growth (Taddei *et al*, 2004), but formed patches at the nuclear membrane upon carbon source exhaustion. We observed the same type of reorganization of nuclear pores, from continuous to patchy, but Nup57 patches and Esc1 patches do not colocalize (**Figure 5C**). At this stage, we cannot conclude whether the reorganization of Esc1 at the nuclear periphery is causally related or just correlated to telomere hyperclustering.

### The mutation of a single serine of Esc1, S1450, recapitulates *esc1Δ* phenotypes regarding telomere clustering during metabolic transitions

Since Esc1 level and localization remains largely unchanged upon quiescence entry, we wondered if Esc1-mediated anchoring could be regulated by a post-translational modification. Esc1 was shown to interact with the H-BRCT domain of the telomere bound protein Sir4, which recognizes a 6 amino-acid domain in Esc1 named the TD-like motif (Faure et al., 2019, Deshpande et al., 2019). An *in vitro* study showed that the phosphorylation of the TD-like domain of Esc1 is critical for its interaction with the Sir4 H-BRCT domain (Deshpande et al., 2019) **(Figure 6A)**. To assess the role of the phosphorylation of Esc1 in the dynamics of telomere clusters during metabolic transitions we built strains mutated for Esc1-S1450. First, to make sure that the Esc1-S1450A protein was stably expressed and correctly localized, we tagged the *esc1-S1450A* mutant with pH-tdGFP in C-terminus using CRISPR-Cas9. As shown in **Figure 6B**, the S1450A mutation does not alter the levels of Esc1 nor its localization during metabolic transitions. Like the WT protein, Esc1-S1450Ap is stable and is localized at the nuclear periphery. Strikingly, the point mutation recapitulates the phenotypes observed in the *esc1Δ* mutant. The *esc1-S1450A* strain forms precocious telomere hyperclusters at 24h (31% for the *esc1-S1450A* vs 7% for WT and 29% for *esc1Δ*) (**Figure 6C, Figure S6**). In stationary phase, the percentage of cells with a hypercluster is strongly increased in the point mutant (80%) compared to the WT (51%) and is similar to that observed in the *esc1Δ* strains (83%). As in *esc1Δ* cells, this high percentage of cells showing telomere hyperclusters reflects the ability of *esc1-S1450A* NQ cells to form hyperclusters (**Figure 6C**, Figure S6).

**Figure 6:**
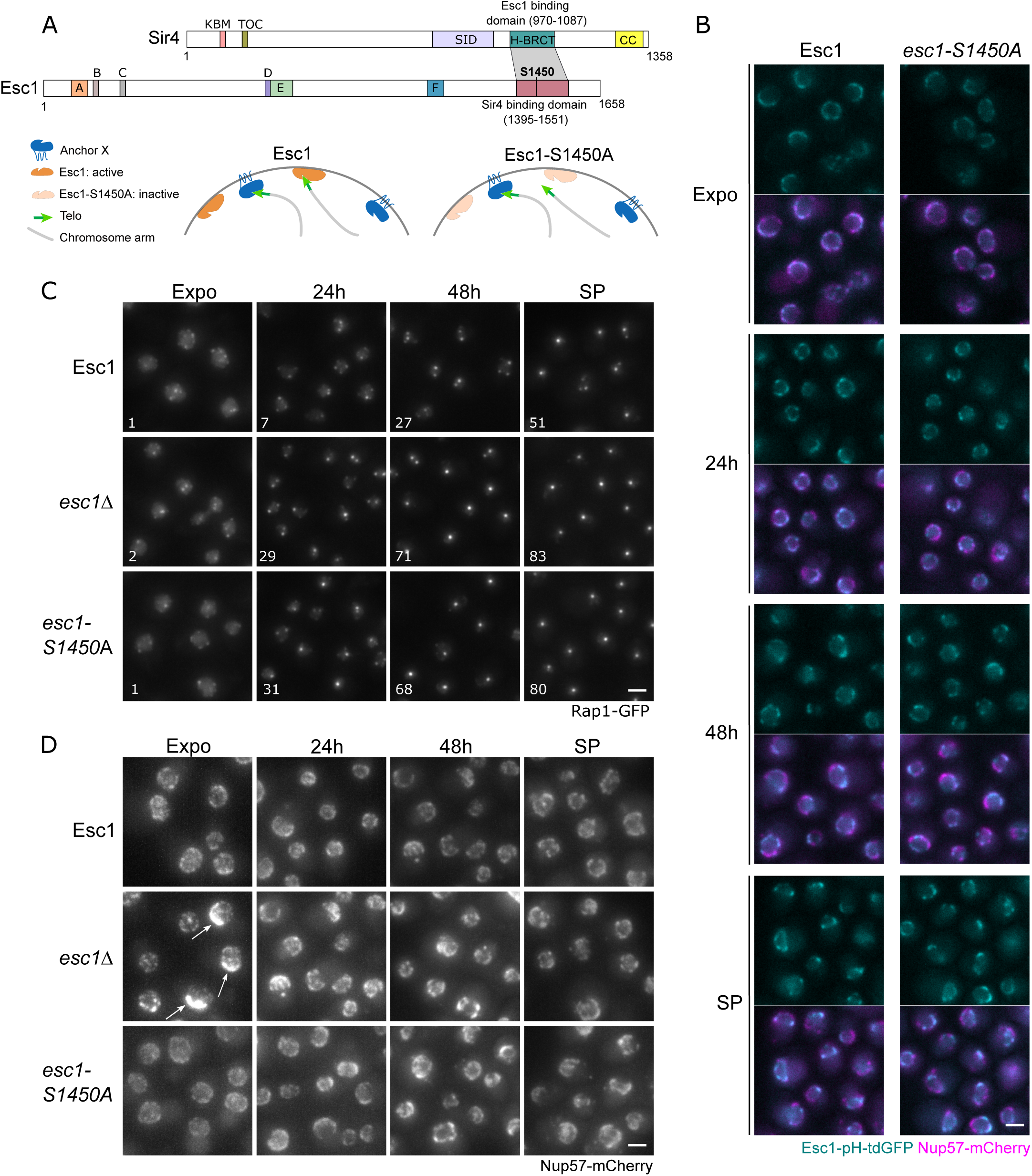
Mutation of Esc1 serine 1450 to alanine phenocopies an *ESC1* deletion. A) Schematic representation of Sir4 and Esc1 conserved domains according to Faure *et al.,* 2019, Deshpande et al., 2019. The grey zone represents the Sir4/Esc1 interaction domain bearing the Serine 1450 of Esc1 (in bold). Schematic representation of telomere anchoring in a WT *ESC1* strain and in the *esc1-S1450A* mutant. B) Representative fluorescent images of strains expressing Nup57-mCherry either with Esc1-pH-tdGFP (yAT4771) or Esc1-S1450A-pH-tdGFP (yAT4285) during exponential phase, at 24h, 48h and during stationary phase. C) Representative fluorescent images of strains expressing Rap1-GFP either WT for *ESC1* (yAT2487) or *esc1Δ* (yAT3811) or bearing the *esc1-S1450A* mutation (yAT4068) during exponential phase, at 24h, 48h and during stationary phase. White numbers at the bottom of each image indicates the percentage of cells with a telomere hypercluster. D) Representative fluorescent images of strains expressing Nup57-mCherry either WT for *ESC1* (yAT4017) or *esc1Δ* (yAT4018) or bearing the *esc1-S1450A* mutation (yAT4286) during exponential phase, at 24h, 48h and during stationary phase. White arrows indicate bright Nup57-mCherry patches. Scale bar is 2µm.

It was previously shown that the deletion of Esc1 impacts the subcellular localization of a subset of nuclear pore proteins, the Sir4-associated Nup complex (among them Nup84, Nup170, Nup157, Nic96, Nup188) without affecting others (Nup53, Nup59). In the *esc1Δ,* those proteins form patches in the nuclear membrane instead of being homogenously distributed in the nuclear envelope (Lapetina *et al*, 2017). To follow the distribution of the nuclear pores in our strains, we used a protein from the inner ring, Nup57 fused to mCherry. We observed that Nup57-mCherry subcellular localization is altered in the *esc1Δ* strain, similarly to the Snup complex proteins (Lapetina *et al*, 2017), forming very bright patches in exponential growing cells in *esc1Δ* (white arrows **Figure 6D**). Interestingly, this is not the case in the Esc1-S1450A point mutant protein that localized to the nuclear envelope like the WT Esc1. Thus, *esc1-S1450A* is a separation-of-function mutant that affects the dynamics of telomere clustering, mimicking *ESC1* deletion phenotypes regarding telomere clustering, without affecting the localization of the Snup complex in the nuclear envelope. In conclusion, serine 1450 is a key residue for Esc1 anchoring function and for the regulation of telomere clustering, possibly through a mechanism involving dynamic phosphorylation of this site during metabolic transitions.

## Discussion

Telomeres undergo spatial reorganization within the nucleus during metabolic transitions (Guidi *et al,* 2015, yet the mechanisms underlying this process remain poorly understood. Since Sir3 is directly involved in telomere clustering in exponential phase (Ruault *et al*, 2011, 2021), we investigated its role in the formation of telomere hyperclusters in quiescent cells. Our study confirmed that Sir3 levels do not increase during metabolic transitions (Guidi *et al*, 2015). Moreover, although we found that Sir3 becomes phosphorylated upon glucose exhaustion, reaching a peak of phosphorylation around 48 hours of culture, this modification is not required for telomere hyperclusters formation; rather it slightly hinders clustering during the transition from exponential to stationary phase. We also observed that Sir3 is sumoylated on lysine 247 during exponential phase and that this modification is strongly reduced after the diauxic shift, coinciding with the appearance of brighter telomere foci. This suggests that Sir3 de-sumoylation promotes clustering. Consistent with this hypothesis, abolishing Sir3 sumoylation leads to a mild increase in telomere clustering throughout the time course to quiescence. Therefore, both phosphorylation and sumoylation of Sir3 appear to counteract telomere clustering.

A more pronounced effect, however, was observed upon deletion of the telomeric anchor Esc1. This is consistent with our physical model of chromosome organization in yeast, which suggests that reducing telomere anchoring at the nuclear periphery promotes hyperclustering. Our modeling reveals that telomere anchoring and hyperclustering counteract each other. One possible explanation is that large telomere clusters cannot form at the nuclear periphery due to limited space for the repulsive steric interactions between chromosomal arms stemming from telomeres. It should be noted that this constraint is mitigated for centromeric clusters, as they do not interact directly with the nuclear envelope or with each other, but are permanently attached to the SPB 200 nm from the nuclear envelope.

Our modeling also shows that hyperclusters cannot form when the interaction between telomeres is too strong. Again, this is due to repulsive interactions between the chromosome arms that prevent the merging of stable clusters. On the other hand, weaker telomere-telomere interaction allows small clusters to dissolve and their individual telomeres to join larger clusters. The size and stability of the largest cluster will increase until a hypercluster is formed. This is analogous to the well-characterized phenomenon Ostwald ripening is phase separation (Ostwald, 1897; Zwicker *et al*, 2025)

This conclusion is apparently in contrast with our previous work showing that increasing Sir3 cellular amount induces the formation of telomere HC, that was initially interpreted as an increase of interaction between telomeres. It is noteworthy that HC induced by Sir3 overexpression also localize away from the nuclear periphery. As we discussed earlier (Ruault *et al*, 2011), Sir3 may shield Sir4 interaction with its perinuclear anchors Mps3 and Esc1 since Esc1, Mps3, and Sir3 all interact with the C-terminal half of Sir4 (Moazed *et al*, 1997; Andrulis *et al*, 2002; Bupp *et al*, 2007). Furthermore, Sir3 competes with Sir4 for binding the telomeric protein Rap1. We thus propose that Sir3 overexpression leads to increased telomere clustering *via* a moderate increase in telomere-telomere interactions and by preventing telomere anchoring.

Our modeling reaches the same conclusion as the one developed for mouse rod cells (Falk *et al*, 2019), namely that losing the perinuclear anchoring of heterochromatin is sufficient to allow hyperclustering of heterochromatin in the nuclear center. However, our finding that hyperclusters cannot form when interactions between heterochromatic loci are too strong was not reported in the model developed by Falk *et al*, 2019. This may be because, in their model, euchromatic regions do not repel each other, thereby preventing the formation of polymer brush observed in our simulations, which is consistent with Hi-C data (Guidi *et al*, 2015; Ruault *et al*, 2021). Another major difference between the two models is the proportion of self-interacting heterochromatin: it represents less than 1% of the yeast genome versus ≈15 % of the simulated mouse genome. Therefore, for all yeast telomeres to cluster within a single hypercluster, the 32 chromosome arms must extend outward from a sphere of 200–300 nm in diameter, whereas heterochromatin in rod cells clusters into a sphere whose diameter is an order of magnitude larger. Finally, in our simulation, telomeres can move along the nuclear envelope, which was not the case for heterochromatin in the mouse model. While this constraint explains the formation of small clusters at the nuclear periphery in the latter case, we also obtain small clusters at the nuclear envelope even in the absence of this constraint. We interpret this phenomenon as a consequence of increased steric interactions at the nuclear envelope due to the geometry of spherically symmetric polymer brushes.

Our genetic analysis confirms that telomere anchoring at the nuclear periphery regulates telomere foci dynamics by counteracting clustering. Based on our genetic and modeling approach, we propose the following model to explain telomere clustering upon metabolic transitions **(Figure 7)**. During exponential growth, telomere foci are anchored to the nuclear periphery through multiple redundant pathways (including Sir4-Esc1, Sir4-Mps3 and Yku-Mps3). When glucose is exhausted, after the diauxic shift, wild-type cells display fewer, brighter telomere foci located at the nuclear periphery. At this stage of the culture (PDS), Esc1 plays a key role in preventing hyperclustering of telomeres – that can occur only away from the nuclear periphery – by maintaining their peripheral anchoring. In the absence of Esc1, telomere hyperclusters form in the center of the nucleus just a few hours after the diauxic shift, whereas in wild-type cells telomeres remain attached to the nuclear periphery forming several clusters in wild-type cells. Thus, telomere anchoring counteracts telomere clustering in PDS cells. At the later stages of the cultures, as cells enter stationary phase, Esc1-dependent anchoring is differentially regulated between quiescent (Q) and non-quiescent (NQ) cells, Esc1-mediated anchoring progressively decreases in Q cells but is maintained in NQ cells. Although HC result mainly from the deactivation of anchoring in Q cells, Sir3 de-sumolyation could also contribute via a moderate increase in telomere-telomere interactions.

**Figure 7.**
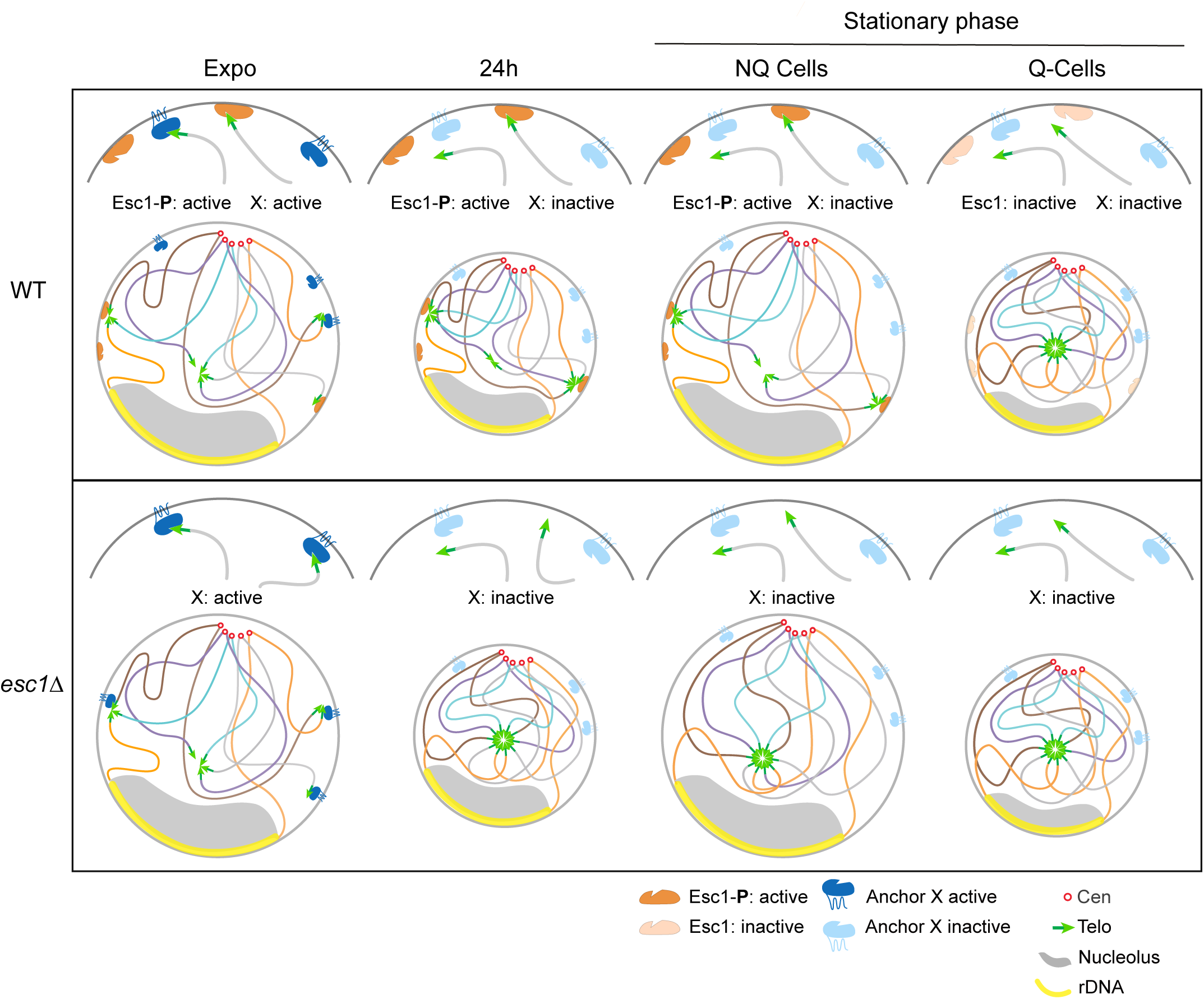

Mechanistically, Esc1 anchors telomeres via a direct interaction between its phosphorylated TD-like domain and the Sir4-HBRCT domain. Indeed, the Sir4-HBRCT domain interacts with a 34 amino acids Esc1 fragment (aa 1440-1473) in the C-terminal domain of Esc1 (Faure *et al*, 2019; Deshpande *et al*, 2019). The Sir4-HBRCT domain also interacts with the Ty5 IN protein (integrase of the Ty5 retrotransposon) through a 6 amino acids motif, the TD domain. Those two Sir4 interacting domains are functionally equivalent since they can substitute for each other (Brady *et al*, 2008). Phosphorylation of the Ty5 IN TD domain is required for its interaction with Sir4 *in vivo* (Dai *et al*, 2007; Brady *et al*, 2008). Similarly, phosphorylation of Esc1-S1450 is required for its interaction with Sir4 *in vitro* (Deshpande *et al*, 2019). We showed that this mutation results in the formation of precocious telomere HC and the formation of HC in NQ cells *in vivo*, indicating that phosphorylation Esc1-S1450 is required for its interaction with Sir4 *in vivo*. Interestingly, Ty5 TD phosphorylation is regulated and reduced under stress conditions, such as starvation from amino acids, nitrogen, or fermentable carbon sources, preventing its interaction with Sir4 (Dai et al., 2007). We propose that Esc1 phosphorylation could similarly be diminished in the absence of carbon sources, specifically in Q cells through a comparable regulatory mechanism. Esc1 dephosphorylation in Q cells may lead to the formation of HC whereas the maintenance of Esc1 phosphorylation in NQ cells could retain telomeres at the nuclear periphery in this cell type. The pathway regulating Ty5p phosphorylation has not been identified yet. Identifying the signaling pathway that regulates Esc1 phosphorylation (either through kinase inactivation or phosphatase activation) will be a crucial step toward understanding how genome organization responds to environmental cues. Interestingly, we showed that the *esc1-S1450A* mutant is a separation-of-function variant that is defective for telomere anchoring but does not alter the organization of nuclear pores. In the future, it could serve as a valuable tool to investigate the functional role of Esc1.

Loss of heterochromatin anchoring has also been documented in metazoans where it is associated with drastic changes in gene expression and replication timing of the affected regions. For instance, upon entry into senescence, Lamin Associated Domains (LADs) undergo a large-scale reorganization and form prominent heterochromatic foci within the nucleoplasm, known as senescence-associated heterochromatic foci, SAHFs (Narita *et al*, 2003; Corpet & Stucki, 2014; Shaban & Gasser, 2025). Another reorganization event, reminiscent of our observations in yeast quiescent cells, occurs during differentiation of rod photoreceptors in nocturnal animals (Solovei *et al*, 2009, 2013). In these cells, during differentiation, chromatin transitions from a conventional to an inverted organization, where heterochromatin clusters in a large compartment in the nuclear interior. As in yeast, this structural transition is slow, one month in mammals, several hours in yeast, probably reflecting difference in genome size.

In mammals and in yeast, heterochromatin anchoring is ensured by multiple redundant pathways. Indeed, in mammals, depletion of LMNB1 alone is not sufficient to induce heterochromatin reorganization during senescence (Sadaie *et al*, 2013), and the formation of inverted nuclei in rod cells requires the inactivation of both LBR and Lamin A/C (Solovei *et al*, 2013). Similarly, we found that in yeast, Esc1 deletion is not sufficient to reorganize telomere clusters in exponentially growing cells, indicating that an Esc1-independent anchoring mechanism is inactivated after the diauxic shift. Overall, heterochromatin anchoring through redundant pathways appears to be conserved across species, a feature which may reflect its functional importance.

Despite our efforts we could not identify the key anchors maintaining telomere at the nuclear periphery during exponential growth in the absence of Esc1. Deleting other known anchors in combination with *ESC1* did not lead to telomere hyperclustering during logarithmic growth, suggesting the existence of additional, yet unidentified, telomeric anchors. Another possible explanation is that telomere clusters are too dynamic during exponential phase to form stable hyperclusters since they are disassembled at each cell division (Laroche *et al*, 2000).

The formation of telomere hyperclusters in the absence of Esc1 after the diauxic shift indicates that the Esc1-independent pathway is inactivated in the absence of glucose. Indeed, abrupt glucose starvation or inactivation of the main glucose sensing pathway, by depleting Cdc35, induces telomere hypercluster formation in the absence of Esc1. We noticed that Cdc35 inactivation induced a stronger response than glucose starvation (16% of telomere hyperclusters at 8h versus 3% at 8h, respectively). These differences may reflect distinct cellular states associated with the two conditions. Indeed, Cdc35 inactivation mimics nutrient deprivation and leads to a G1 arrest (Matsumoto *et al*, 1983; Werner-Washburne & Singer, 1993), whereas abrupt glucose starvation causes cell cycle arrest at various stages (Wood *et al*, 2020). In agreement with this hypothesis, prolonged G1 alpha-factor arrest induces a moderate increase of telomere clustering in *esc1Δ* cells. We thus conclude that Esc1-independent anchoring involves at least two distinct anchoring mechanims, one regulated by the cell cycle (inactivated in G1) and the other by glucose signaling (inactived in the absence of glucose).

We have demonstrated that, in *S. cerevisiae*, telomere anchoring plays a key role in orchestrating telomere clustering in response to metabolism. The dynamic relocalization of telomeres during metabolic transitions, and its differential behavior in quiescent (Q) versus non-quiescent (NQ) cells, suggest that telomere spatial organization is not a passive feature of nuclear architecture. Rather, it may play an active role in coordinating genome functions with cellular physiology, like the lamina in metazoans. Spatial control of telomeres may contribute directly to the fine-tuning of gene expression near chromosome ends which are enriched for genes that play key roles in the organism’s interaction with its environment (Hocher & Taddei, 2020). Alternatively, it could be important to maintain telomere integrity under nutrient limitation. We recently showed that intergenic accumulation of RNA PolII during quiescence facilitates a swift transcriptional restart upon exit from this state (Baquero Pérez *et al*, 2025). In a similar way, telomere hyperclustering might serve as a regulatory or protective mechanism helping cells to ensure a coordinated metabolic response and/or preserve genome stability when cells withstand harsh environmental conditions.

## Methods

### Growth conditions

All the strains used in this study are described in Supplemental Table S1. Yeast cells were grown at 30°C in YPD (1% yeast extract, 2% bactopeptone and 2% glucose) autoclaved for 15 minutes at 110°C. For induction of quiescence by carbon source exhaustion, yeast cells were grown overnight in YPD, diluted at 0.1 OD/ml in the morning in aerated flask (using Hirschmann™ SILICOSEN™ silicone caps) and grown with agitation (250 rpm) for several days: 24h, 48h, 3 days and stationary phase time-points being collected directly from the flask. Cells reached an OD_600nm_ around 50-60 ODs/ml after 5 to 7 days of culture. To isolate Quiescent (Q) and non-quiescent (NQ) yeast cells from stationary phase, cells were loaded on a density gradient; the high-density (HD) and low-density fractions (LD), respectively enriched for Q and NQ cells, were recovered as described in Baquero-Perez *et al,* 2025. For starvation experiment from exponential phase, cells were grown to OD_600nm_ = 1, pelleted and resuspended in water or YP medium in an aerated flask which was placed at 30°C with agitation. Levels of glucose in the culture were assayed using glucose test strips from Precision laboratories; test strips were dipped in the culture for 2 seconds, the color was read 3 minutes later. For Cdc35-AID-9myc degradation, cells were grown to early exponential phase (OD_600nm_ = 0.4 OD/ml) and IAA was added directly to the culture at a final concentration of 500mM. To arrest the cells in the G1 phase, alpha-factor was added directly to exponential phase cultures (OD_600nm_ = 0.4 OD/ml) to a final concentration of 5 µM.

### Strain constructions

Gene deletions and gene tagging were performed by PCR-based gene targeting as previously described (Longtine *et al*, 1998; Janke *et al*, 2004) or using the CRISPR-Cas9 technologies (DiCarlo *et al*, 2013; Soreanu *et al*, 2018). To overexpress *SIR3* in the BY strain, we targeted an integrative plasmid containing the *SIR3* gene under the control of its own promoter (pAT349) at the *his3Δ0* locus, the number of plasmid insertions was evaluated by measuring the level of the Sir3 protein by western blot.

### Protein immunoblotting

For total protein extracts, 1.2 × 10^8^ cells were harvested and precipitated on ice for 10 min in 10 mls of 10% trichloroacetic acid (TCA). Cells were pelleted by centrifugation, resuspended in a 200 µl volume of 10% TCA and glass beads were added to the meniscus. Cells were lysed for 10 minutes in a bead bitter. Extracts were centrifuged at 4°C at 10000 rpm for 5 minutes and the pellets resuspended in 100 µl of TCA-sample buffer (50 mM Tris-HCl pH 6.8, 100 mM dithiothreitol, 2% SDS, 0.1% bromophenol blue, and 10% glycerol containing 200 mM of unbuffered Tris solution). The extracted proteins were denatured 5 min at 95°C and loaded (10 µl of total extracts per well) on a 7.5% mini-Protean TGX Stain-Free Precast gel from Biorad. Proteins were transferred on a nitrocellulose membrane which was blocked in 1xPBS, 5% milk, 0.2% Tween (blocking buffer) for 30 min and then incubated overnight with the anti-Sir3 antibody at 1/5000 in blocking buffer (Ruault *et al*, 2011). The blots were washed 3 times 10 min in blocking buffer and incubated for 45 minutes with an anti-rabbit HRP at 1/10000 (Jackson ImmunoResearch Laboratories) in blocking buffer. The blots were then washed 3 times in 1xPBS, 5% milk (washing buffer) and 5 minutes in 1xPBS before revelation. Normalization of the loading was done using the stain-free technology from Biorad. For phosphatase treatment (calf intestine alkaline phosphatase – CIP – from Roche), samples were diluted 1:2 in a final volume of 20 µl with phosphatase buffer (1X), 0,5mM MgSO4, 20mM Tris pH8,8 and incubated for 3 hours at 37°C with 10U of CIP or without CIP for undigested control.

### Microscopy

Images were acquired on a wide-field microscopy system controlled by MetaMorph software (Molecular Devices), based on an inverted microscope (Nikon TE2000) equipped with a 100×/1.4NA immersion objective. Images were captured using a Cmos camera (ORCA-flash C11440, Hamamatsu, Japan), and a Spectra X light engine lamp (Lumencor, Inc, Beaverton, OR, USA). The images shown are maximum intensity projections of z-stacks acquired with a 200 nm step. For each experiment, all images are acquired using the same acquisition settings.

### Telo-FISH

The probe, containing 4.8 kb of Y’ element and TG repeats (pEL42H10; Louis & Borts, 1995) was obtained by PCR amplification using primer pair oAT151-GAAGAATTGGCCTGCTCTTG /oAT152-CCGTAAGCTCGTCAATTATT. The PCR product was purified using the Qiaquick PCR purification kit from Qiagen. 1 µg of the purified product was labeled using the nick translation kit from Jena Bioscience (Atto488 NT Labeling Kit); the labeling reaction was performed at 15°C for 90 min; the labeled DNA was purified using the Qiaquick PCR purification kit from Qiagen and eluted in 30 µl of water; 100 µl of formamide, 50 µl of 40% dextran sulfate, 20 µl of 20x SSC were added to the probe for a final concentration of 50% formamide, 10% dextran sulfate, 2X SSC. The FISH protocol we used is adapted from Laenen *et al*. (in preparation). 20 OD of cells (1 OD corresponding to 10^7^ cells) were grown to mid–logarithmic phase (1–2 × 10^7^ cells/ml) and harvested at 1,200g for 5 min at RT. Cells were fixed in 20 ml of 4% paraformaldehyde for 15 min at 30°C, quenched by adding 125 mM Glycine, spun down for 5 min at 800g and washed with 10 mls of 125 mM Glycine for 5 minutes. After centrifugation for 5 min at 800g, cells were washed once in 10 mls of 1x PBS, centrifuged again and resuspended in 1.5 mls of 0.1 M EDTA-KOH pH 8.0, 10 mM DTT for 10 min at 30°C with gentle agitation. Cells were then collected at 800g and the pellet was carefully resuspended in 1.25 ml YPD - 1.2 M sorbitol. Next, cells were spheroplasted at 30°C with Zymolyase (60 mg/ml Zymolyase-100T to 1 ml YPD-sorbitol cell suspension) for 10 min for the exponential cultures and 30 minutes for the 24h cultures. Spheroplasting was stopped by the addition of 40 ml YPD - 1.2 M sorbitol. Cells were washed twice in YPD - 1.2 M sorbitol, and the pellet was resuspended in 500 µl of 1x PBS. Cells were next attached on a glass coverslip coated with Poly-L-Lysine and air dry in a small petri dish. The coverslips were washed with 1x PBS, 0.5% Triton X-100 at RT (15 min while shaking) and incubated in 2 mL of 1x PBS, 20% glycerol at RT for 2 hours. Next, the coverslips in their buffer were frozen using liquid nitrogen and thawed at RT, 4 times successively. They were then washed in 1x PBS, 0.05% Triton X- 100 for 5 min, rinsed in water, incubated in 0.1N HCl at RT for 5 min and washed in 1x PBS, 0.05% Triton X-100 for 5 min. The coverslips were prehybridized in 2x SSC / 50% formamide for 1 day at RT in the dark. 20 µl of hybridization buffer (2x SSC /50% formamide /10% dextran sulfate) containing the Y’ probe (1/4 dilution) was added to a microscope slide; the coverslip was gently settled on the drop of probe solution and fixed on the slide using fixogum. The microscope slide was placed on a heating block at 80°C from 3 minutes to denature DNA and incubated overnight in a humid chamber at 37°C in the dark. The next day, the coverslips were washed twice with 0.2x SSC, 0.2% Tween-20 at 45°C for 7 min. Then, they were briefly washed; first in 4x SSC, 0.2% Tween-20 and then three times in 2x SSC. The coverslips were post-fixed in 4% paraformaldehyde, 1x PBS for 10 min, washed twice in 1x PBS for 10 min, and once in 2x SSC. Finally, the nucleus was stained with DAPI; the coverslips were incubated in a 2x SSC, 1,25 µg/ml DAPI for 5 min and then washed twice in 2x SSC before imaging.

### Image quantifications

Quantification of Rap1-GFP foci was performed using an in-house algorithm, NuFoQ, described in Baquero Pérez *et al*, 2025. Briefly, all 3D stacks have been deconvolved using SVI Huygens (Ponti *et al*, 2007) with the Classical Maximum Likelihood Estimation algorithm at 20 iterations and a quality change stopping criterion of 0.05. Based on deconvolved images, NuFoQ (https://github.com/mgPICT/NuFoQ) fully segments spots in 3D and extracts their intensities as total integrated pixel values from raw images. Rap1-GFP total nuclear signal is taken as nuclear staining. Cluster (focus) intensities are calculated relative to nuclear staining. A hypercluster is defined as a cluster with an intensity at least 4 times above the average intensity of clusters from the wild-type in exponential phase. Proportions in pie charts represent the percentage of cells without focus, 1 focus, 2 foci, 3 foci and 4 or more foci. Measurement of the distances between the telomere clusters and the nuclear periphery were obtained with a modified version of NuFoQ: NuFoQal (https://github.com/mgPICT/NuFoQal). Briefly, the raw image of the tagged nucleoporin and the background illumination of the Rap1-GFP signal show respectively the outer rim and the inside of the nuclei. A morphological opening on the GFP signal allows for removal of the foci. Both images are then merged to generate a nuclei image. On this new image, the yeast nuclei are first detected in 2D with a top hat filtered sum projection using the pretrained cyto2 model of Cellpose (Stringer *et al*, 2021; Pachitariu & Stringer, 2022). The detected candidates are separated with a Voronoi tesselation and then fully segmented in 3D independently from each other using fuzzy C-means and keeping the 3D convex hull of the resulting pixel classification. Foci are identified using the same methodology than NufoQ.

Cell size was obtained from light transmitted images using MIC-MAQ (Microscopy Images of Cells - Multi Analyses and Quantifications; https://github.com/MultimodalImagingCenter/MIC-MAQ), a Fiji plugin dedicated to automatic segmentation of nuclei and/or cells for quantifications. The automatic segmentation was performed using Cellpose’s cyto2 model (Stringer *et al*, 2021).

### Computational model

The simulations employ a bead model of chromosomes, with 850 bp per bead for all loci except rDNA, that is modeled as 853 larger beads, 1600 bp per bead. There are three telomeric beads at each end of the chromosome. Interactions between loci are governed by a Lennard-Jones (LJ) potential:

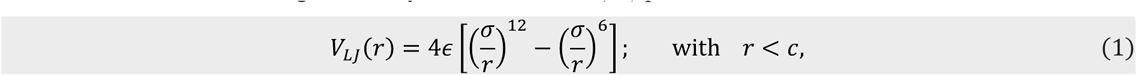

where *σ* = 1 defines the LJ unit, *c* is the cutoff, and *ε* is the interaction energy. Bonds between loci along a chromsome are modeled with a quadratic-quartic potential:

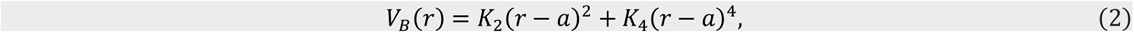

where *a* is the bond length.

The interaction between the nuclear envelope and the loci is given by a LJ potential as in Bystricky *et al*, 2004, in function of the distance of the locus from the nuclear envelope. The LJ potential between the nuclear envelope and non-telomeric loci, as well as between non-telomeric loci in the chromosomal arms, is made purely repulsive by setting the cutoff *c* = *σ*. The LJ potential between the nuclear envelope and nucleolus loci is attractive with energy *∈*_*NE*_ = 10. To achieve proper rDNA segregation on the nuclear envelope, rDNA beads are larger than normal with a radius *σ* = 2. The LJ potential between the nuclear envelope and rDNA beads is attractive with energy *∈*_*NE*_ = 10. The LJ interaction energies *∈*_*TT*_ and *∈*_*TE*_, which describe the interaction strengths between telomeres, and between a telomere and the nuclear envelope, respectively can be referred as the wetting and the condensation parameter. Temperature is maintained at T = 1 using a Langevin-based thermostat. For the first 5 million steps, *∈*_*TT*_ is set to zero, together with the attractive tail of *∈*_*TE*_. The strength of the rDNA interaction is turned on first, and the nucleolus allowed to migrate toward the nuclear envelope, so as to avoid unphysical entanglement with the remaining chromsomes. Most simulations didn’t reach a steady state within 30 million timesteps. The simulation is performed in LJ units, the nuclear radius is set to 30 LJ. The calibration of the LJ unit to nm as well as the timesteps to seconds is described in the following section. Each simulation was run at least 10 times, each with a different random number seed. All simulations are run using the LAMMPS molecular dynamics software (Thompson *et al*, 2022). Simulation configuration parameters are uploaded as sample runs in Supplementary Material 1, allowing a reproducibility of the numerical result.

#### Validation and calibration in the computational model

The model is validated and LJ units are fitted to the physical system with two steps. First, the LJ unit is determined by extracting the 3D distance scaling in function of the genomic distance variation *r*(*s*) along the chromosome and adjusting the LJ unit to fit measured values the two-loci imaging data from (Bystricky *et al*, 2004; Arbona *et al*, 2017) for different loci (Figure S2B). The two-loci distance scaling in these experiments is compatible to the random walk model, *r*(*s*) ∝ *s*^1/2^. The fitted parameter allows us to determine that the 30 LJ units used for the nuclear radius correspond to 0.75 µm, compatible to experiments. Second, the simulation timestep parameter is scaled to fit the Mean Squared Displacement (MSD) *r*^2^(*t*) of single loci tracking *in vivo*. For this, we use the single particle tracking datasets in the thousands of milliseconds ranges from (Hajjoul *et al*, 2013; Arbona *et al*, 2017; Miné-Hattab *et al*, 2017). The single locus MSD scaling in these experiments is compatible to the Rouse model, *r*^2^(*t*) ∝ *t*^1/2^ . In those datasets the dynamics of telomeres differ significantly from those for central loci and those differences are not fully captured by our model (Figure S2C). We decided to focus solely on the dynamics for the telomere 3R to obtain a single fitting time unit. We determine that 100,000 timesteps corresponds to 25 seconds, and a full 30 million timestep simulation represents 7.5 10^3^ seconds (∼ 125 minutes) of real time.

### Numerical analysis, definition of a cluster

A cluster is defined as a group of telomeres, each of which is within 1.5 LJ units of another as in Ten Wolde & Frenkel, 1998. For a given timestep in the simulation, the positions of all telomeres are scanned pairwise and assigned to a cluster if the proximity condition is satisfied. We further restrict our cluster definition to those containing two or more telomeres (six or more telomeric beads in our model). We can then study the evolution of cluster spatial distributions and cluster sizes as a function of time.

## Acknowledgments

The authors thank the members of the Taddei laboratory and UMR3664 for helpful discussions. The A.T. team was financially supported by funding from Agence Nationale pour la Recherche DeSynLE (ANR-22-CE120013- 01). The authors greatly acknowledge the PICT-IBiSA@Pasteur Imaging Facility of the Institut Curie, a member of the France Bioimaging National Infrastructure (ANR-24-INBS-0005 FBI BIOGEN). We thank Patricia Le Baccon, head of the microscopy platform for her technical support and assistance, Cédric Messaoudi for sharing the MIC-MAQ Fiji plugin with us. This work used the Expanse computer nodes at the San Diego Supercomputer Center through allocation BIO230022 from the Advanced Cyberinfrastructure Coordination Ecosystem: Services & Support (ACCESS) program, which is supported by U.S. National Science Foundation grants #2138259, #2138286, #2138307, #2137403, and #2138296. The National Science Foundation and the Mayent-Rothschild Sabbatical program at Institut Curie supported the work of K. B. B. The National Science Foundation had no role in study design, data collection and analysis, decision to publish, or preparation of the manuscript. The views presented here are not those of the National Science Foundation and represent solely the views of the authors. V.F.S., B.D.K. also acknowledge travel support from the CNRS “Genome Architecture and Function 2023” in Sofia (Bulgaria) that was a key meeting for discussions over the project.

## Author contributions

M.R., I.L., M.B.P. generated strains and performed experiments, A.E. carried out FISH experiments. J.E.H. and D.W. generated and characterized the Cdc35 degron strain. M.G. created NuFoQ and NuFoQal. M.R. analyzed the microscopy experiments. V.F.S., K.B.B. and B.C. performed the modeling. A.T. and M.R. contributed to the design of the experiments. A.T., V.F.S and M.R. contributed to the interpretation of the data, the drafting of the figures and the writing of the manuscript. A.T. and K.B.B. obtained funding.

---

**Figure S1:**
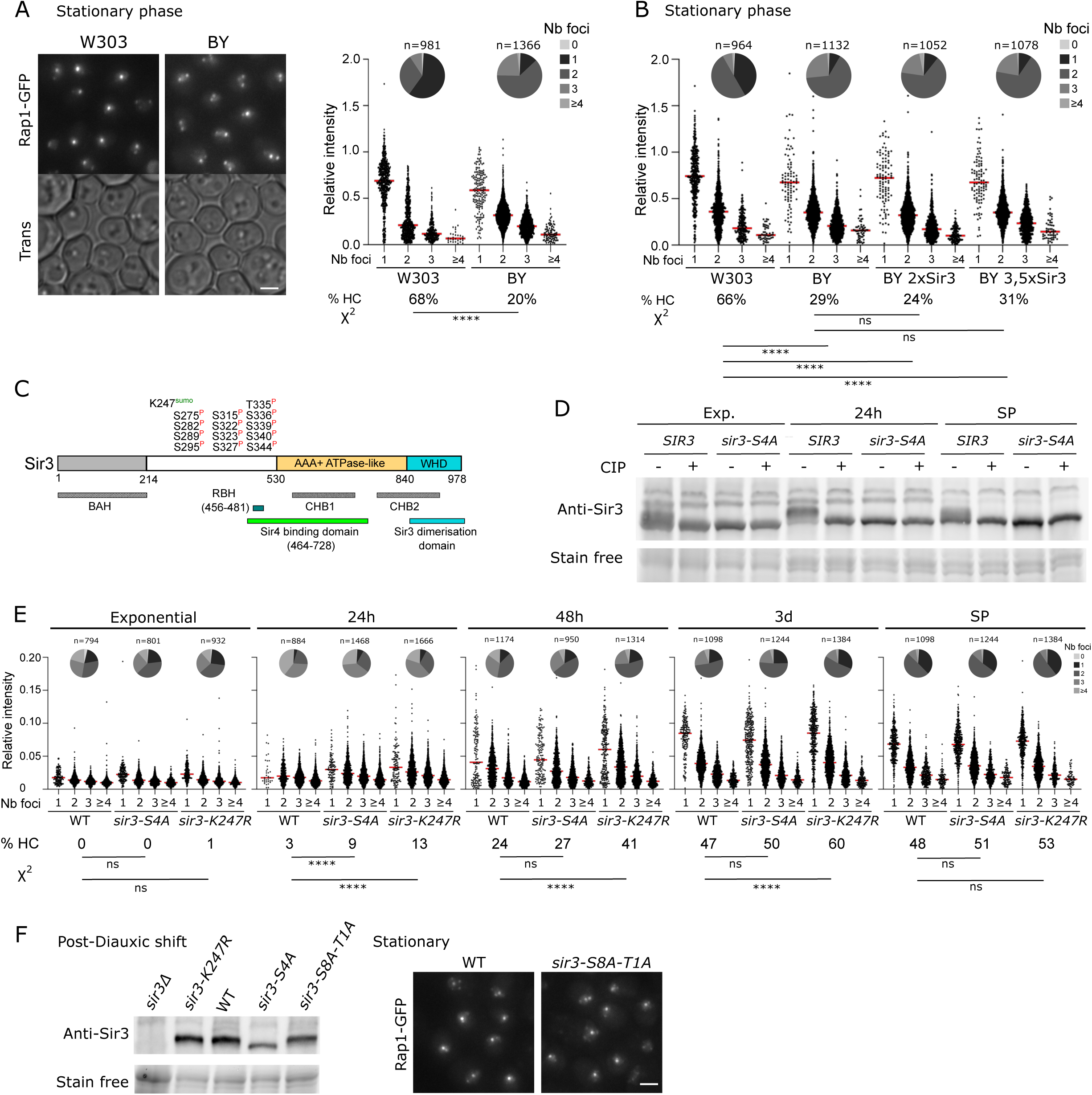
A) Comparison of telomere clustering in W303 and BY4741 backgrounds during stationary phase. Left: Representative fluorescent and transmitted light images of Rap1-GFP strains during stationary phase - WT W303 (yAT2487) and WT BY4741 (yAT1950). Scale bar is 2µm. Right: Quantifications of telomere clustering. Dot plots represent the cluster intensity relative to the total nuclear signal in cells with 1, 2, 3 or 4 and more foci (red bar is the median). The percentage of cells with a hypercluster is indicated for each strain at the bottom of the graph. Statistical analysis was done using a two-sided Chi-square test with a confidence interval of 99%; **** = p < 0.0001. The pie charts represent the distribution of the cells according to the number of foci per cell, n is the total number of cells analyzed. B) Related to Figure 1C: Quantifications of relative intensities, number of clusters per cells and percentage of cells with a hypercluster as in Figure S1A. C) Schematic representation of Sir3 domains and annotations of Sir3 modifications (Sumoylation in green, phosphorylation in red). BAH: Bromo Adjacent Homology domain. CHD1 and CHD2: C-terminal Histone binding Domain 1 and 2. D) Immunoblots using the Sir3 antibody against crude extracts, with or without CIP (phosphatase) treatment, in the WT (yAT2487) and in the *sir3-S4A* mutant during exponential phase, at 24h and 7 days. E) Related to Figure 1G: Quantifications of relative intensities, number of clusters per cells and percentage of cells with a hypercluster as in Figure S1A. F) Left: Immunoblot using the Sir3 antibody on crude extracts from sir3Δ (yAT2553), *sir3-K247R* (yAT3868), WT (yAT2487), *sir3-S4A* (yAT3905) and *sir3-S8A-T1A* (yAT3906) mutants after the diauxic shift. Right: Representative fluorescent images of Rap1-GFP strains - WT (yAT2487) and *sir3-S8A-T1A* (yAT3906) - during stationary phase. Scale bar is 2µm.

**Figure S2:**
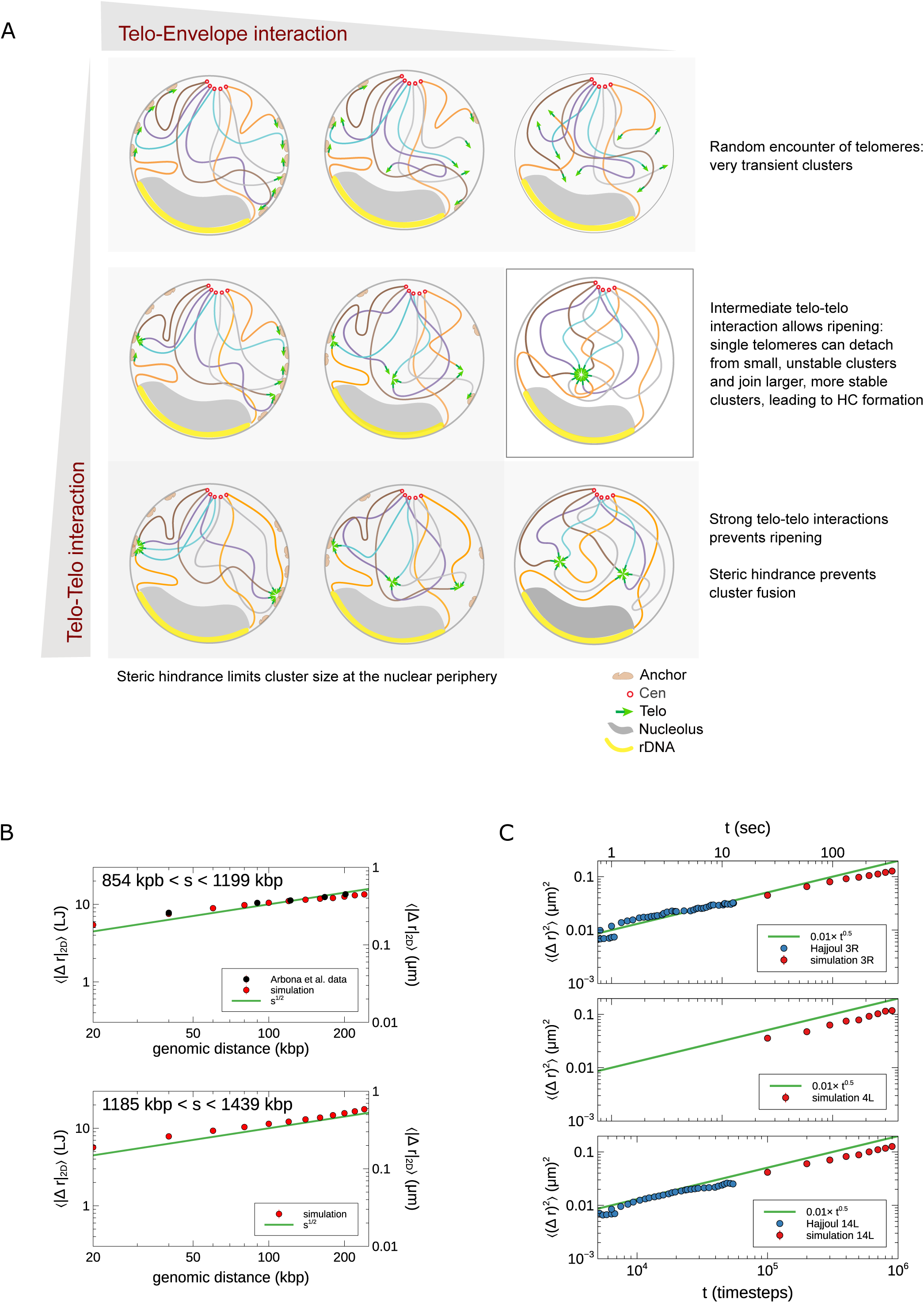
A) Model representing the impact of Telomere-Envelope interaction and Telomere-Telomere interaction on telomere clustering. B) Scaling of the two-loci distance in function of genomic distance, for two regions on chromosome 4. Comparison of experimental data (Arbona *et al,* 2017), fitted on the same data, and simulations. C) Scaling of the mean square displacement of a single locus in function of time, for telomeres 3R, 4L and 14L (Hajjoul *et al*, 2013). Comparison of experimental data, fitted on the dataset for telomere 3R, and simulations.

**Figure S3:**
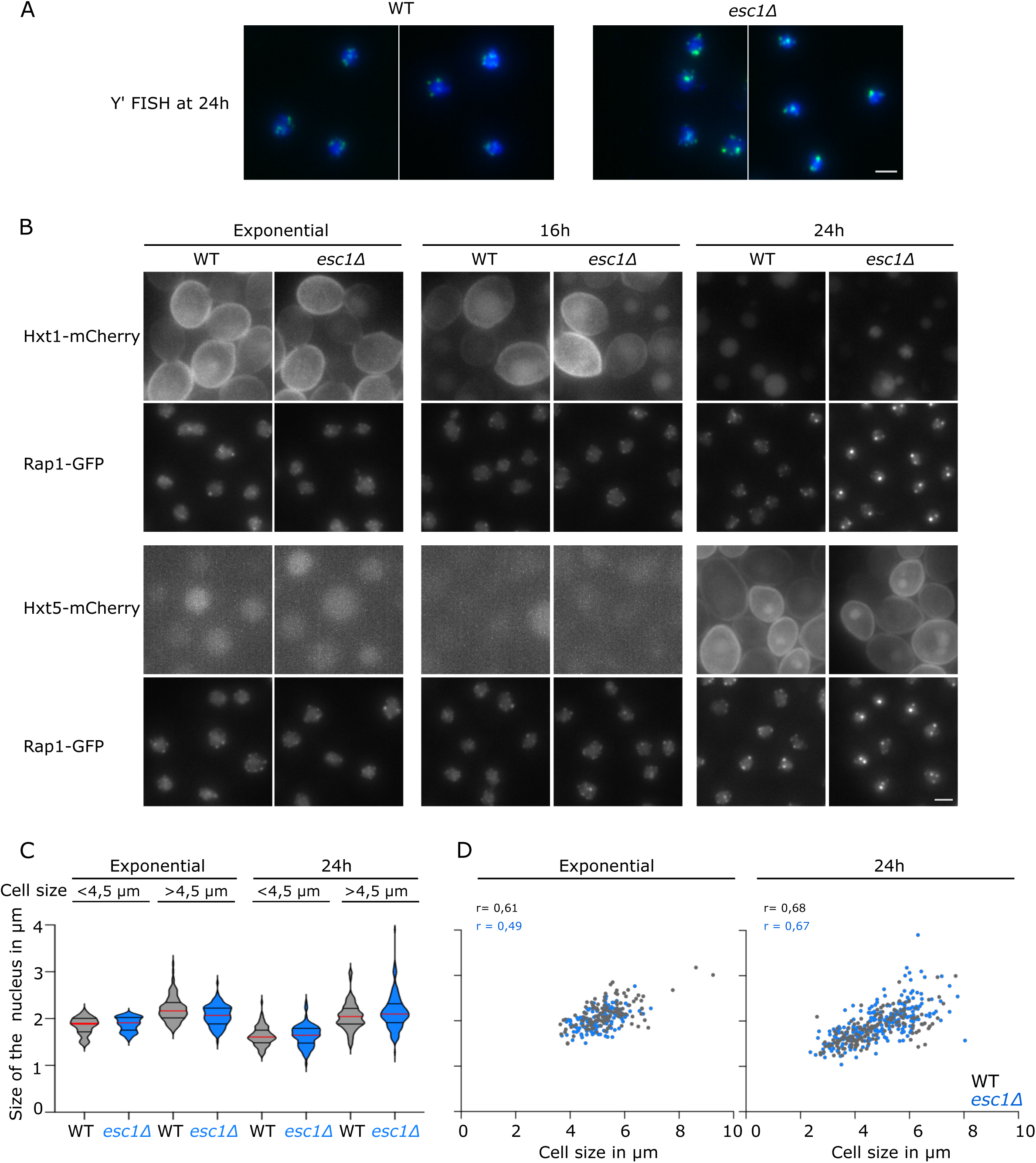
A) Representative images of Y’FISH experiments (green signal) in WT (yAT2487) and *esc1Δ* (yAT3811) strains at 24h of culture. Nuclei are labelled using DAPI staining (blue signal). Scale bar is 2µm. B) Representative fluorescent images of Rap1-GFP and glucose transporters tagged with mCherry, either Hxt1 (WT-yAT3900 and *esc1Δ*-yAT4404) or Hxt5 (WT-yAT3899 and *esc1Δ*-yAT4410) were acquired during exponential growth, at 16h and 24h. Scale bar is 2µm. Hxt1, a low-affinity glucose transporter, is expressed when glucose levels are high and localized at the plasma membrane in those conditions; when glucose levels are low, *HXT1* is repressed and degraded. Hxt5, a high-affinity glucose transporter, is expressed only prior glucose depletion and localized at the plasma membrane. C) Violin plot showing the size of the nucleus in small cells (Feret’s diameter < 4.5 µm) and in larger cells (diameter ≥ 4.5µm) in WT (yAT4017) and *esc1Δ* (yAT4018) strains during exponential phase and 24h. D) Scatter plot with the data from C) showing the correlation between the nuclear and cell size. Pearson correlation coefficient r is indicated for each set of data.

**Figure S4:**
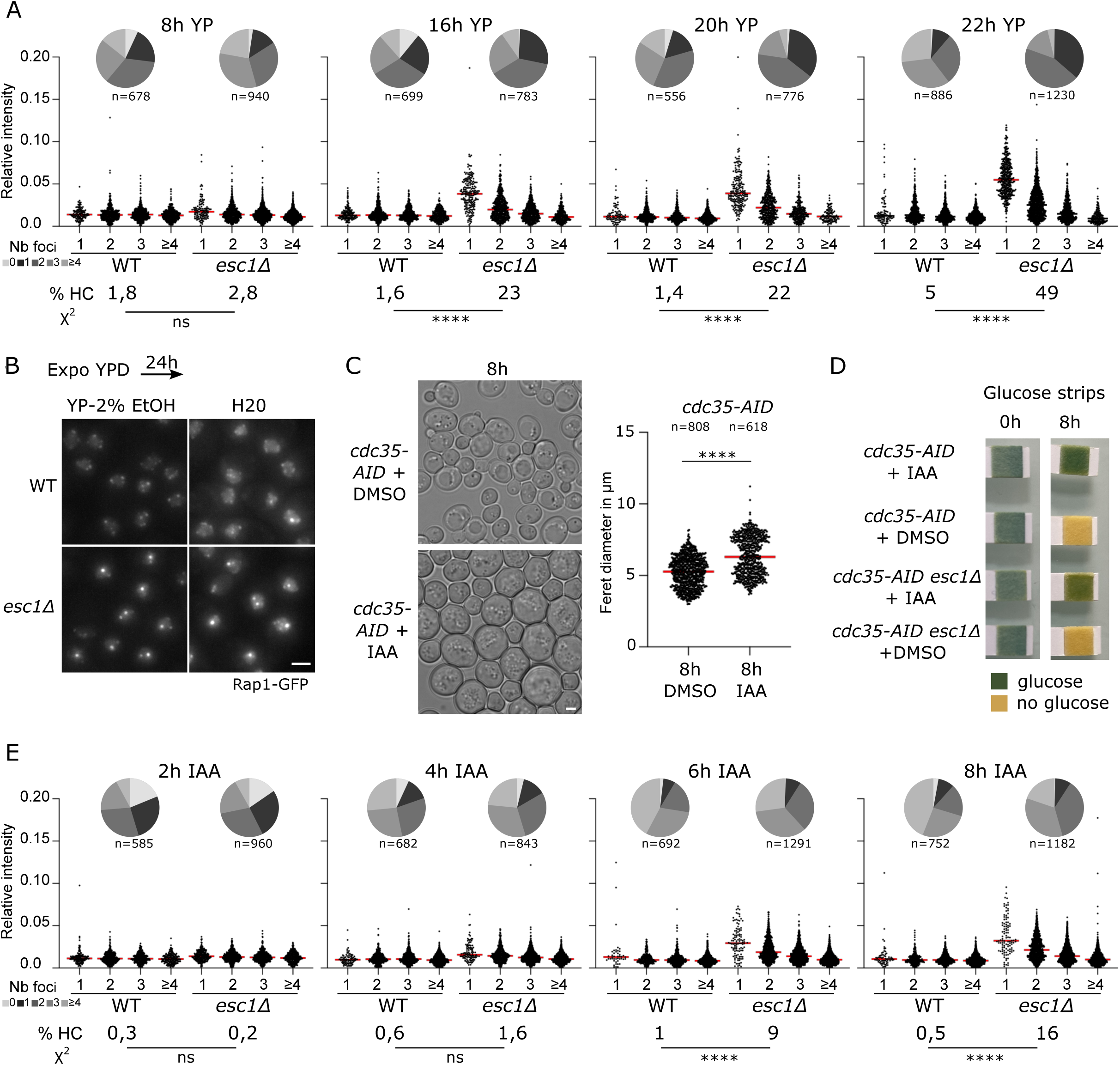
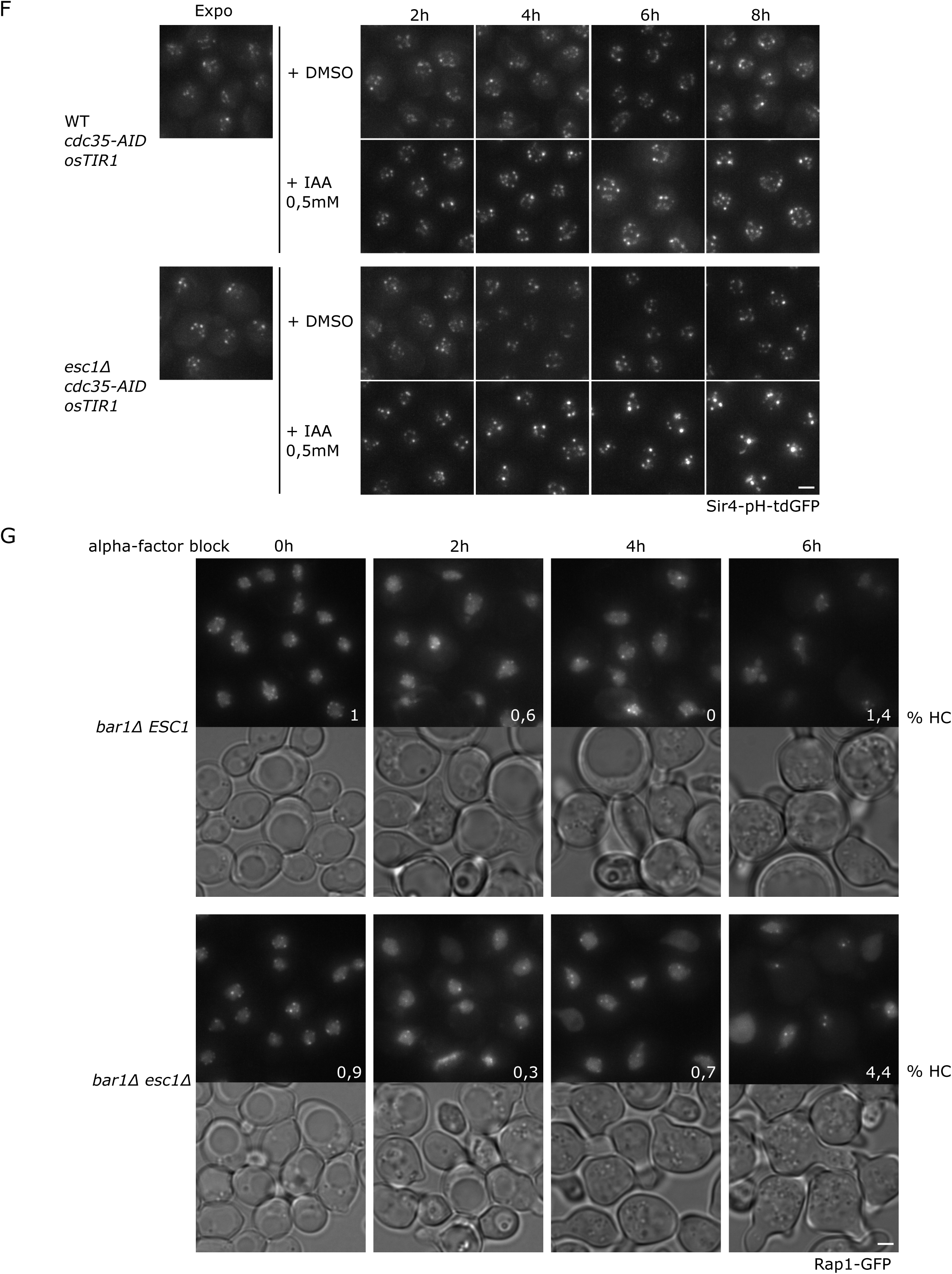
A) Quantifications of telomere clustering related to Figure 4B: Dot plots represent the cluster intensity relative to the total nuclear signal in cells with 1, 2, 3 or 4 and more foci (red bar is the median). The pie charts represent the distribution of the cells according to the number of foci per cell, n is the total number of cells analyzed. The percentage of cells with a hypercluster is indicated below the graphs. Statistical analysis was done using a two-sided Chi-square test with a confidence interval of 99%; **** = p < 0.0001. B) Representative fluorescent images of WT (yAT2487) and *esc1Δ* (yAT3811) strains. Cells were grown to exponential phase (OD_600nm_ =1) and switched to YP with 2% Ethanol or water for 24h before imaging. C) Representative light transmitted images of the WT strain (yAT4827) used in Figure S4C (8h time point with DMSO or IAA). Dot plots represent the cell size (Feret’s diameter) measured in these two samples. Red bar is the median. n is the number of cells analyzed. Statistical analysis was done using a two-sided Welch’s t test with a confidence interval of 99%; **** = p < 0.0001. D) Glucose strips indicating the presence (Green) or the absence (Yellow) of glucose in the medium of the following strains *CDC35-AID pGPD-TIR1 SIR4-pH-tdGFP* (yAT4827) and *esc1Δ CDC35-AID pGPD-TIR1 SIR4-pH-tdGFP* (yAT4842) grown in exponential phase at 0h and after 8h of DMSO or IAA treatment. E) Quantifications of telomere clustering related to Figure 4C: Dot plots represent the cluster intensity relative to the total nuclear signal in cells with 1, 2, 3 or 4 and more foci (red bar: median intensity). The pie charts represent the distribution of the cells according to the number of foci per cell, n is the total number of cells analyzed. The percentage of cells with a hypercluster is indicated below the graphs. Statistical analysis was done using a two-sided Chi-square test with a confidence interval of 99%; **** = p < 0.0001. F) Representative fluorescent images of strains expressing Cdc35-AID, Tir1p under the *GPD* promoter, Sir4-pHtdGFP with or without *ESC1* (yAT4827 and yAT4842, respectively). Cells were grown to exponential phase and treated with DMSO or IAA (0,5mM) at time 0. Images were taken every 2h from 0 to 8h. G) Representative fluorescent images of the telomere-associated protein Rap1 tagged with GFP in *bar1Δ* strains and their corresponding light transmitted images. WT *ESC1* (yAT4752) and *esc1Δ* (yAT4753) strains were grown to exponential phase and α-factor was added to the culture at a final concentration of 5µM. Images were taken at 0h, 2h, 4h and 6h of treatment. White numbers on the fluorescent images represent the percentage of cells with a hypercluster. Scale bar is 2µm.

**Figure S5:**
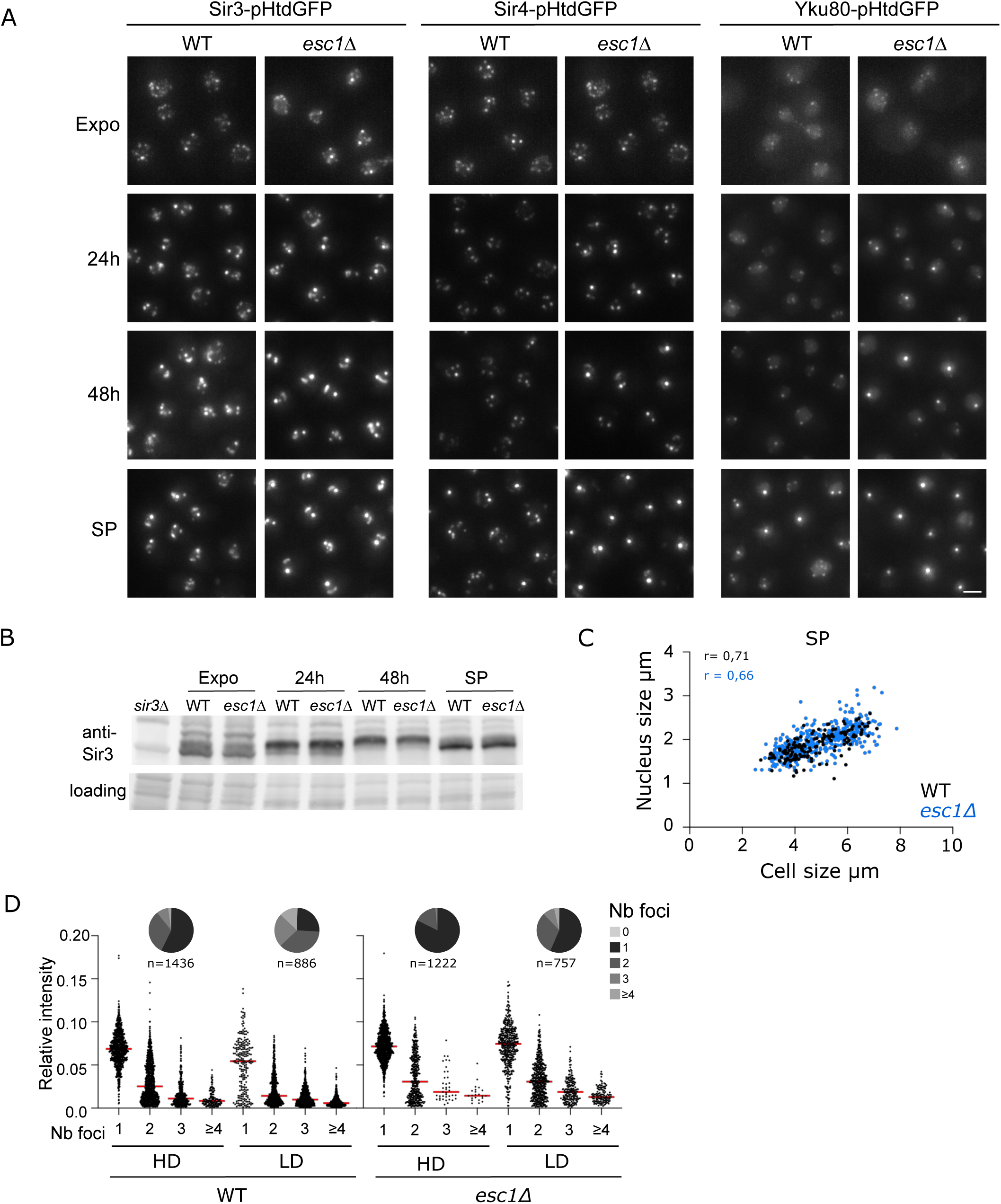
A) Representative fluorescent images of strains expressing Sir3-pHtdGFP, Sir4-pHtdGFP and Yku80-pHtdGFP in the presence or the absence of *ESC1*. Images of WT *SIR3-pHtdGFP* (yAT4155), *esc1Δ SIR3-pHtdGFP* (yAT4196), WT *SIR4-pHtdGFP* (yAT4156), *esc1Δ SIR4-pHtdGFP* (yAT4197), WT *YKU80-pHtdGFP* (yAT4356), *esc1Δ YKU80-pHtdGFP* (yAT4466) were acquired during exponential phase, at 24h, at 48h and during stationary phase. Scale bar is 2µm. B) Immunoblots using the Sir3 antibody. Crude extracts were obtained from the WT (yAT2487) and *esc1Δ* strains (yAT3811) during exponential phase, at 24h, at 48h and during stationary phase. Control loading is shown below the immunoblot using the strain-free technology from Biorad. C) Violin plot showing the size of the nucleus in small cells (Feret’s diameter < 4.5 µm) and in larger cells (diameter ≥ 4.5µm) in WT (yAT4017) and *esc1Δ* (yAT4018) strains during exponential phase and 24h. D) Scatter plot obtained with the data from C) showing the correlation between the nuclear and cell size. Pearson correlation coefficient r is indicated for each set of data. D) Quantifications of telomere clustering related to Figure 5B: Dot plots represent the cluster intensity relative to the total nuclear signal in cells with 1, 2, 3 or 4 and more foci (red bar is the median). The pie charts represent the distribution of the cells according to the number of foci per cell, n is the total number of cells analyzed.

**Figure S6:**
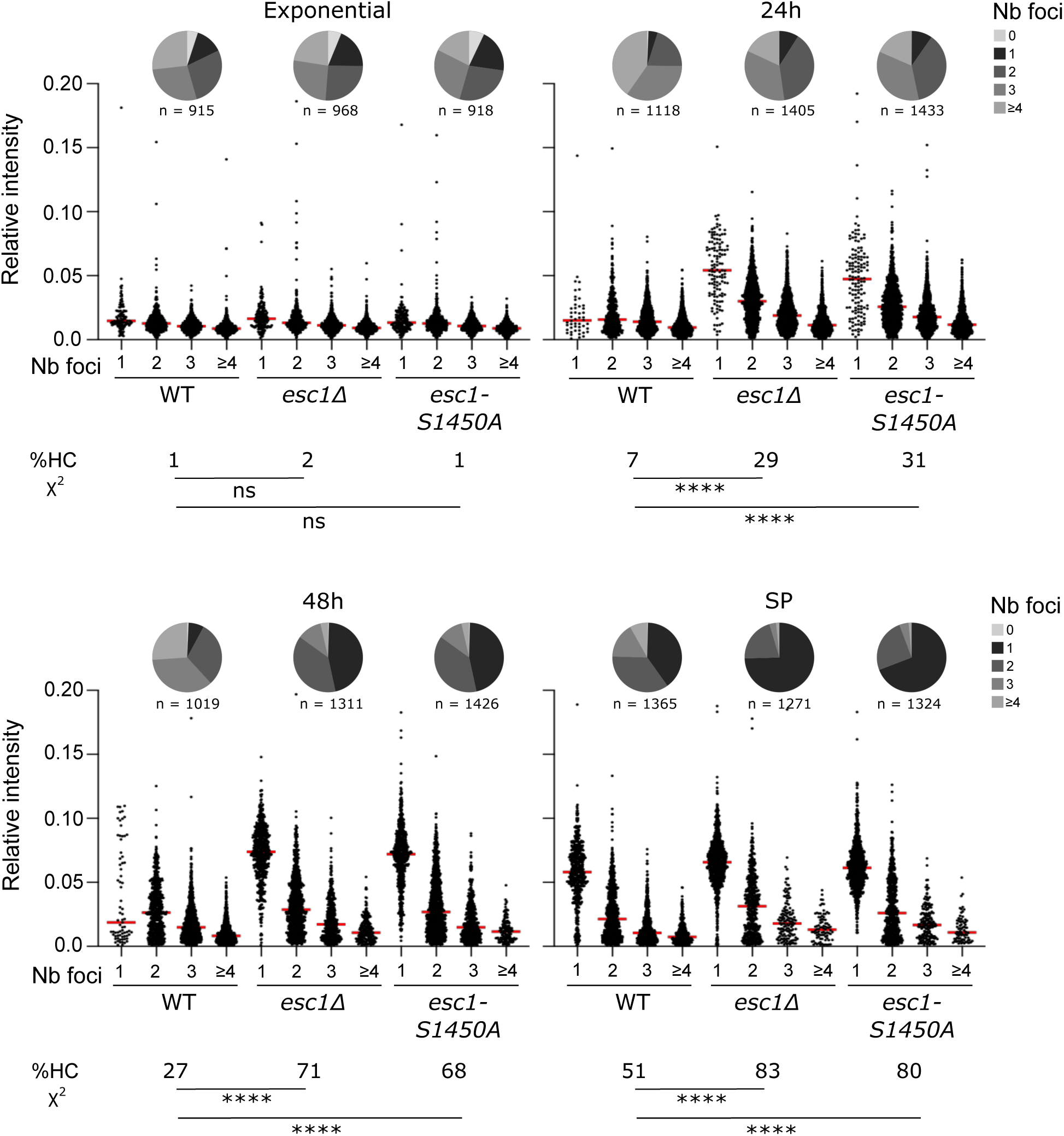
Quantifications of telomere clustering related to Figure 6C: Dot plots represent the cluster intensity relative to the total nuclear signal in cells with 1, 2, 3 or 4 and more foci (red bar is the median). The percentage of cells with hyperclusters is indicated below each plot. Statistical analysis was done using a two-sided Chi-square test with a confidence interval of 99%; **** = p < 0.0001. The pie charts represent the distribution of the cells according to the number of foci per cell, n is the total number of cells analyzed.

**Supplemental Table 1:**
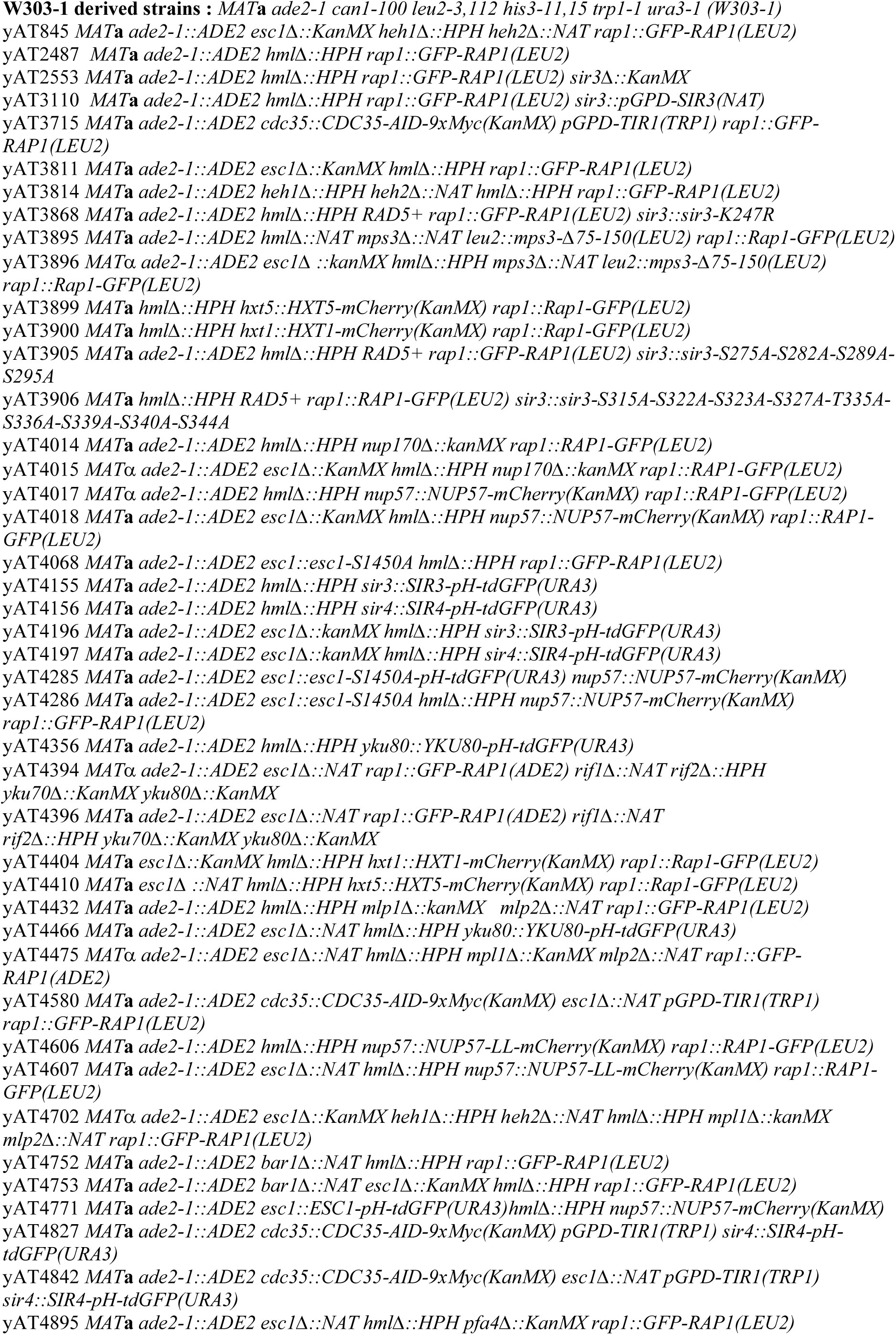
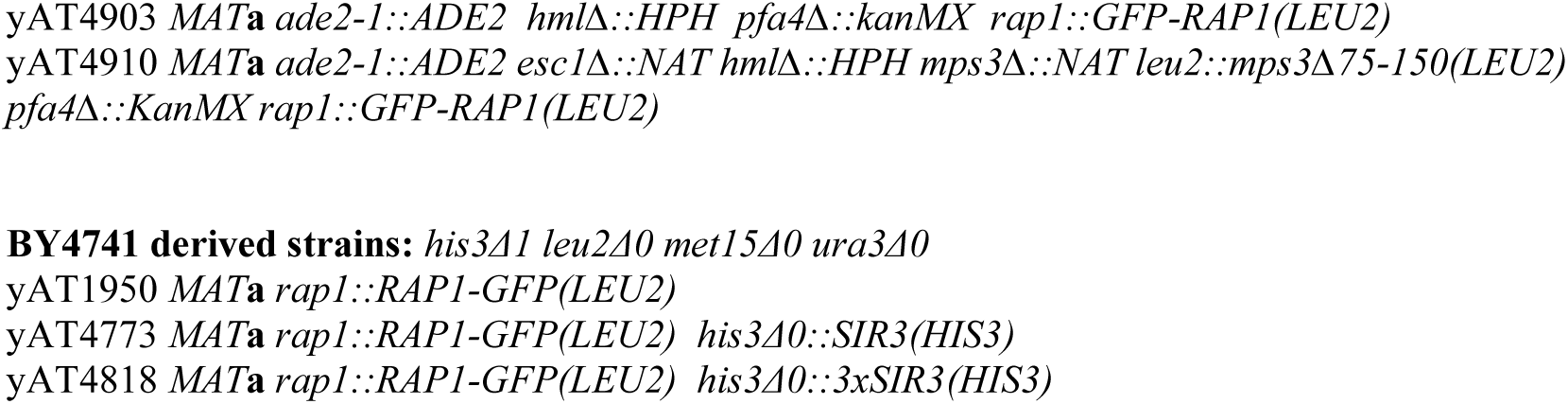
List of the strains used in the study.

