## Supplementary Material 1 for "Esc1-mediated anchoring regulates telomere clustering in response to metabolic changes": Makefile

```
log.lammps	: Y1.lmp Body Header interactions softinteractions variables
	pwd; lmprun Y1.lmp
Bconfig.dat	: CNXconfig.dat
	CNXinit_9.0; cat Header Body > Bconfig.dat
Y2		: Y2.lmp
	pwd; lmprun Y2.lmp
Y3		: Y3.lmp
	pwd; lmprun Y3.lmp
Y4		: Y4.lmp
	pwd; lmprun Y4.lmp
Y5		: Y5.lmp
	pwd; lmprun Y5.lmp
Y6		: Y6.lmp
	pwd; lmprun Y6.lmp
Y7		: Y7.lmp
	pwd; lmprun Y7.lmp
Y8		: Y8.lmp
	pwd; lmprun Y8.lmp
Y9		: Y9.lmp
	pwd; lmprun Y9.lmp
realclean	:	clean
	rm -f Bconfig.dat Body Header Atom*
clean		:	purgeout
	rm -f atom*.dat *.jpg *.avi DropletSize* */DropletSize* restart_state.* log.* *~ slurm* beadtrack* *.pdf r*.dat zone*.dat droplet*.dat
purgeout	:
	rm -f timeheat.dat radialheat.dat focusheat.dat zonemaxout.dat zone*out.dat *balance.dat time*.dat r2ds*.dat rsqt*.dat atom_t.dat atom_t_full.dat atom_c.dat
```
