## Supplementary Material 1 for "Esc1-mediated anchoring regulates telomere clustering in response to metabolic changes": Cluster_Histogram.pdf

ID: Nucleo HX3_3.0_1.7_0.5_1.6
wall potential: lj126
staged rDNA
$W_{rDNA}$=2.0; rDNA scale: 16
$\sigma_{rDNA}=W_{rDNA}$
$c_{rDNA}=1.25\ \sigma_{rDNA}$
$\varepsilon_{RE}=5$; $\sigma_{RE}=\sigma_{rDNA}$
$c_{RE}=1.40\ \sigma_{rDNA}$
$\varepsilon_{NE}=10$
$\varepsilon_{TE}=0.5$; $\sigma_{TE}=1.0$; $c_{TE}=2.5$
$\varepsilon_{TT}=1.6$; $\sigma_{TT}=1.0$ $c_{TT}=1.2$
$\varepsilon_{n2}=0.0$
timesteps: 30M
combined runs: 10
$\varepsilon_{TE}$ turned on at t=5M

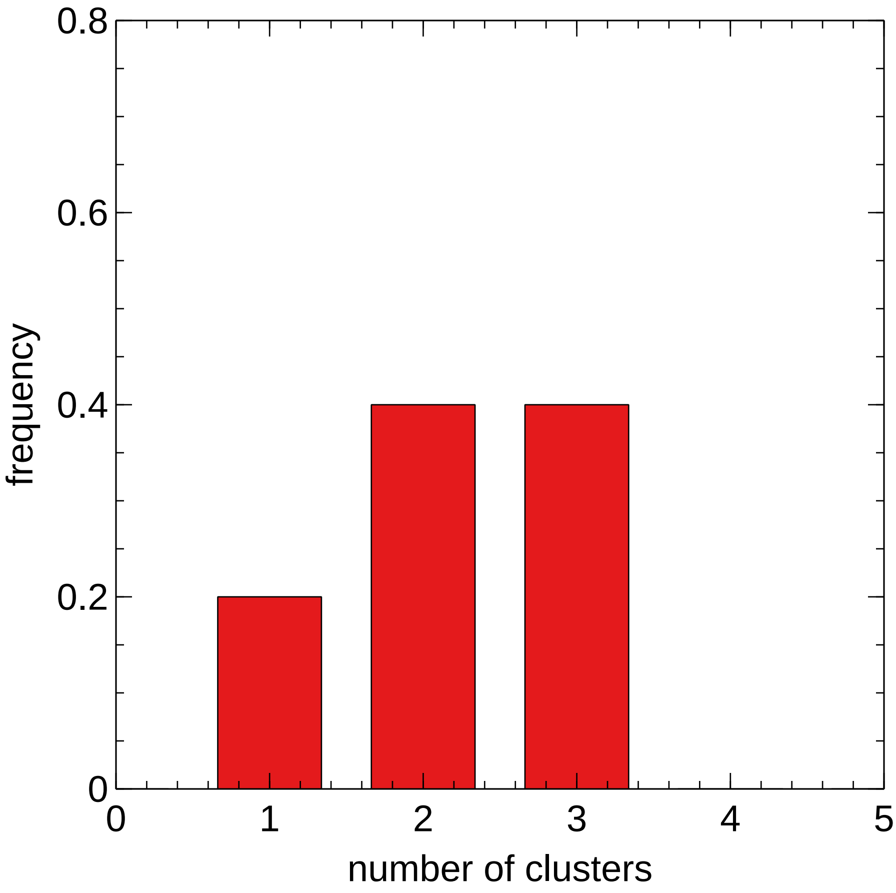
