## Supplementary figures and images for "Esc1-mediated anchoring regulates telomere clustering in response to metabolic changes"

### FocusHeat.pdf

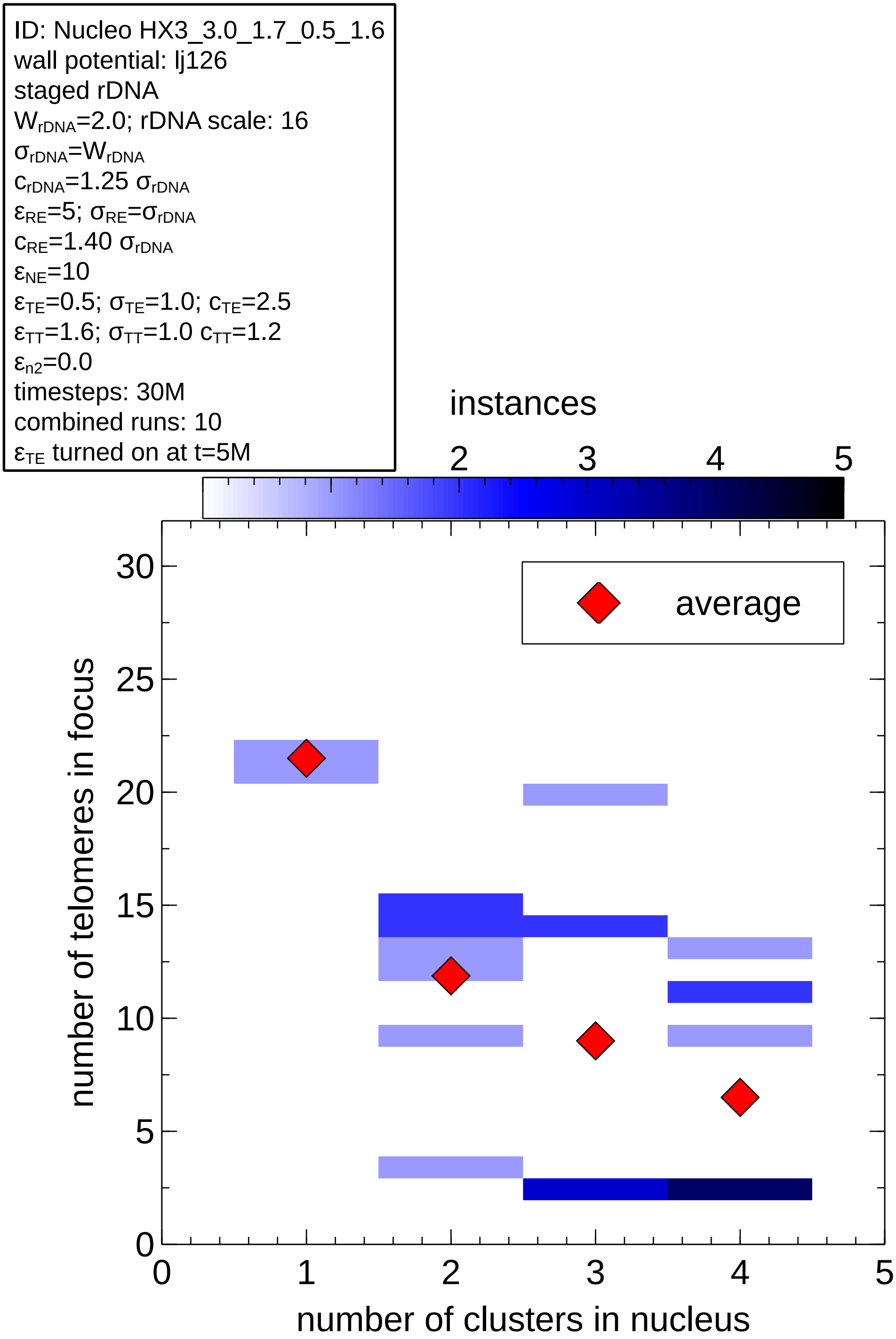

### Heat_Radial-no_label.pdf

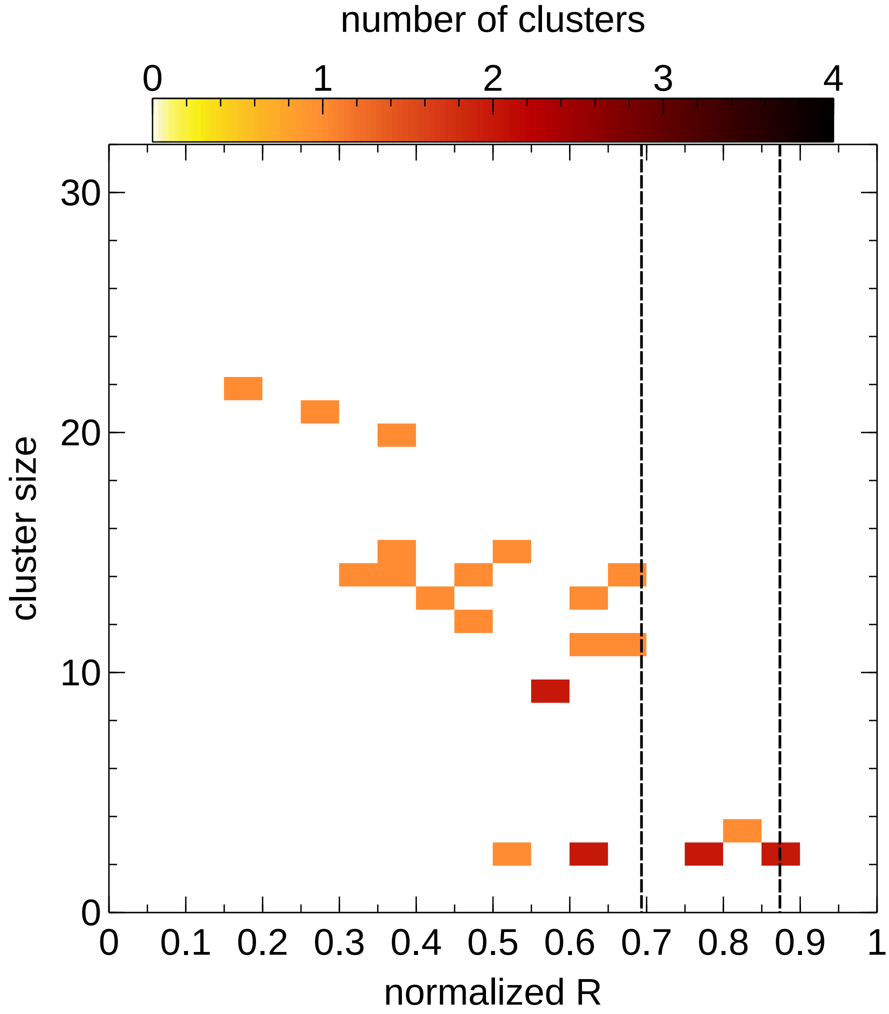

### Heat_Radial.pdf

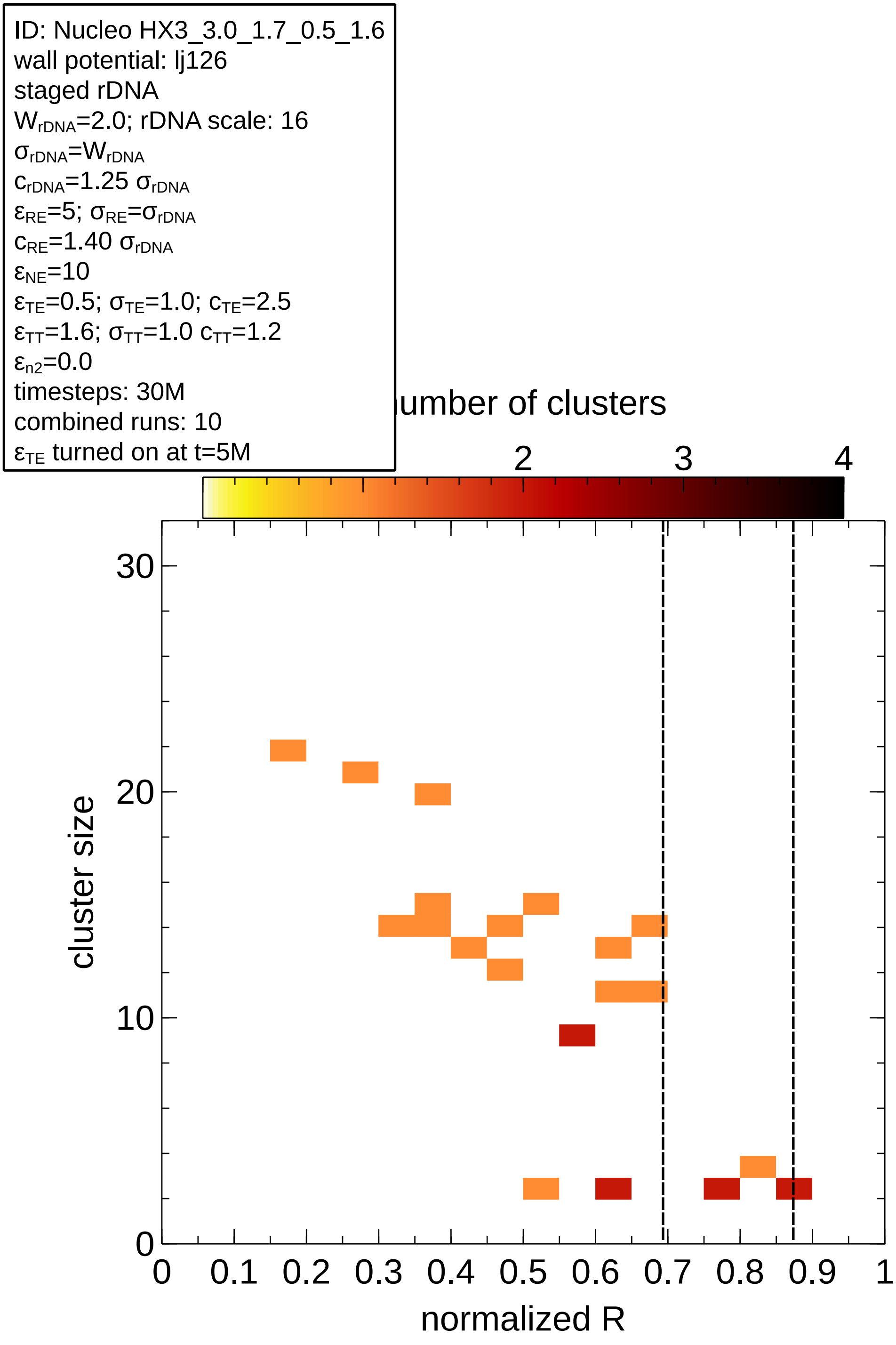

### Heat_Time.pdf

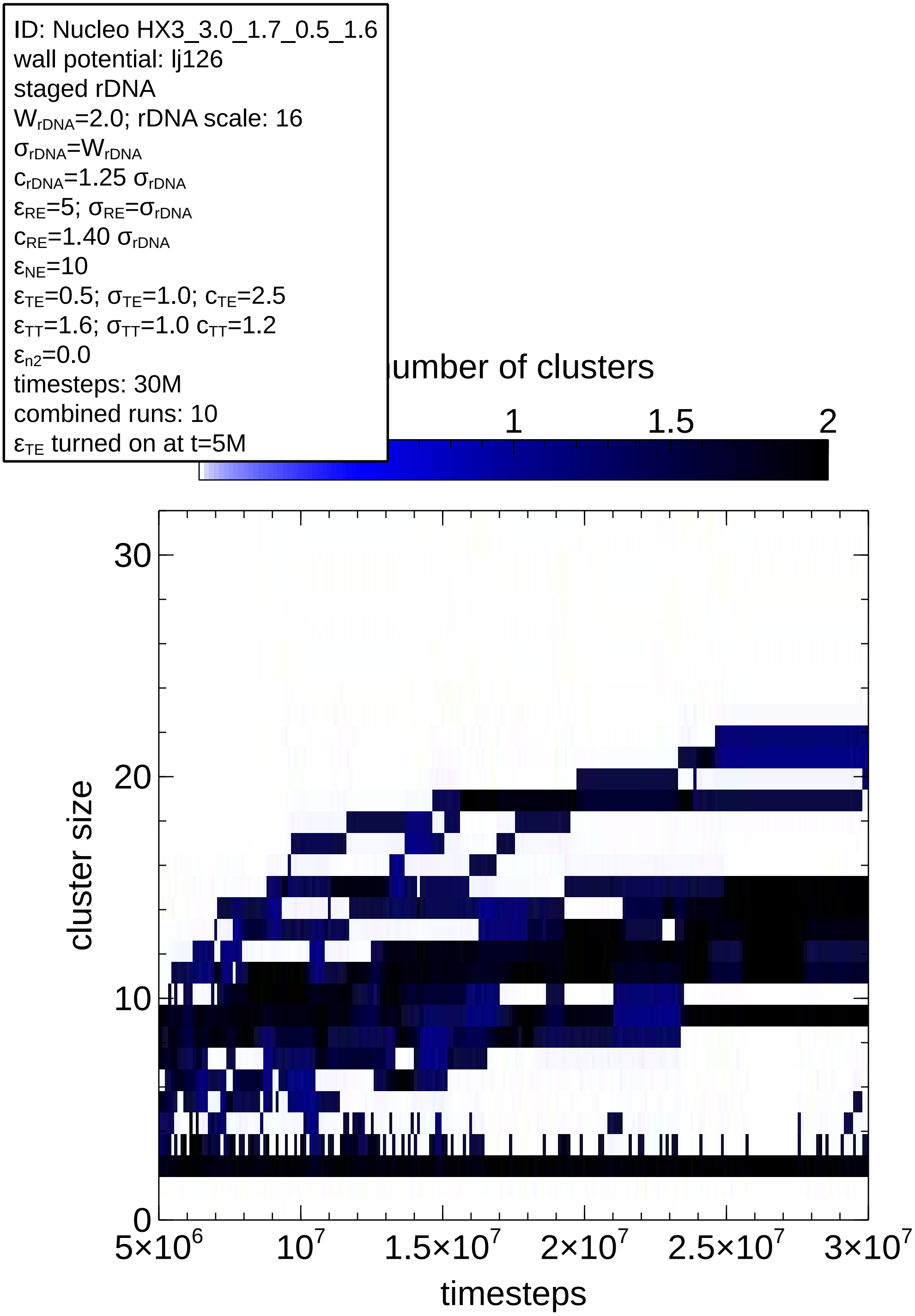

### MaxStack.jpg

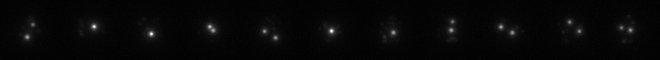

### Pie.pdf

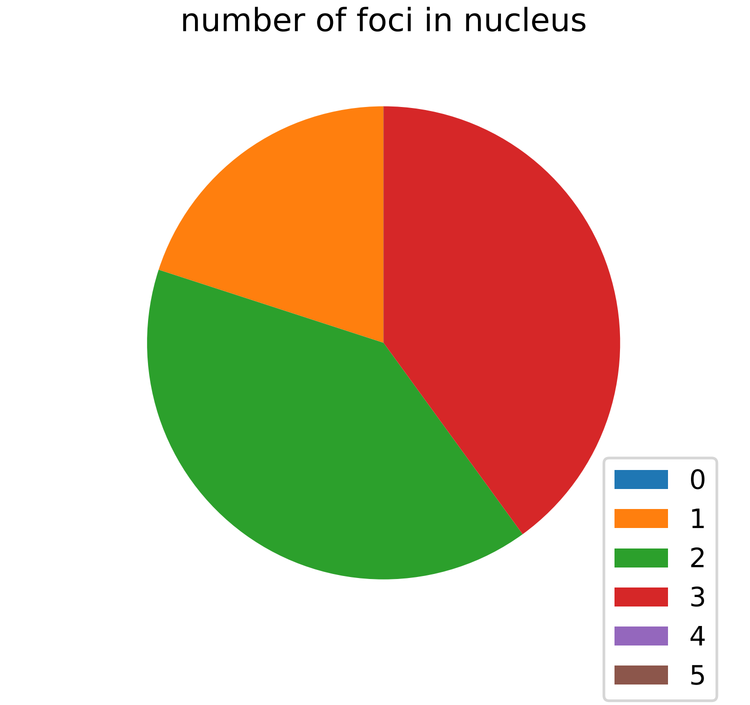

### R2DS.pdf

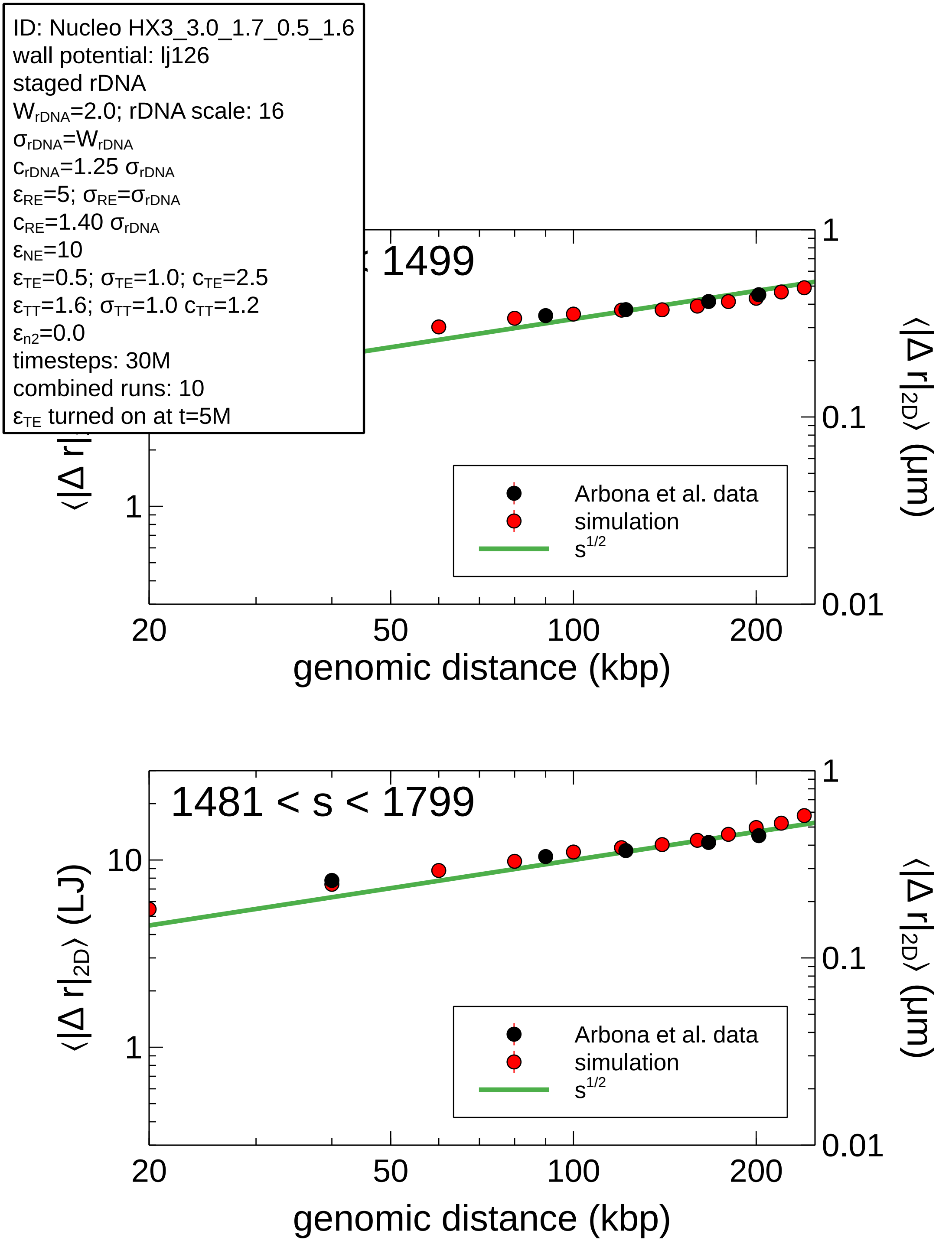

### Reslice of TabStack.jpg

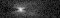
